# The Impact of Glyphosate-Based Herbicides and Their Components on *Daphnia Magna*

**DOI:** 10.1101/794156

**Authors:** Katherine Duan, Alexander Kish, Leanna Kish, Peter Faletra, Kelly Salmon

## Abstract

Recent studies suggest glyphosate-based herbicides (GBHs) are more harmful to animals than suggested by the EPA and industry-funded studies. Both glyphosate and the only known “other” ingredient in GBHs, polyethoxylated tallow amine (POEA), have been implicated as safety hazards. In this study, we investigated the effects of the commercial GBHs Roundup^®^, Rodeo^®^ and the two known GBH ingredients, POEA and glyphosate, on the survival and heart rates of *Daphnia magna*. *D. magna* were exposed to the retail herbicide mixture and the individual components dissolved in water to mimic possible environmental exposure. When exposed to Roundup^®^ and Rodeo^®^, *D. magna* survival and heart rates declined following a dose-response pattern. A commercial formulation of Roundup^®^ containing 98% unlisted ingredients had the greatest effect on heart and survival rates, followed by two formulations of Rodeo with 4.62% unlisted ingredients and 1.72% unlisted ingredients, respectively. The Rodeo^®^ formulation with 1.72% unlisted ingredients had an equal concentration of glyphosate as the Roundup^®^ formulation, suggesting that the negative effects of GBHs are influenced by the unlisted ingredients. Although differences in survival rates were not observed between controls and glyphosate groups, groups exposed to glyphosate alone generally showed a significant (p<0.05) effect on *D. magna* heart rates. Heart rates following POEA exposure were consistently and, in most cases, significantly (p<0.05) lower than controls. POEA caused a decrease in survival rate for all concentrations, but followed a dose-response pattern only in the three highest concentrations. A Mock-GBH, made with POEA and glyphosate, significantly (p<0.05) lowered heart rates at some higher concentrations, with no dose-response pattern. The Mock-GBH negatively affected survival rates at approximately the same level as POEA alone. The heart rate data suggest that there are undisclosed ingredients in Roundup^®^ and Rodeo^®^ other than POEA and glyphosate that negatively affect *D. magna* since glyphosate and POEA combined yielded less pronounced negative responses than the full GBH products.

## 1 Introduction

Although N-(Phosphonomethyl)glycine (glyphosate) was first synthesized in 1950, its efficacy as an herbicide was not reported until the early 1970s (1,2). Glyphosate was initially patented by Monsanto in 1971 (US 3799758) and, by the mid-1970s, marketed with the trade name Roundup^®^. In this herbicide formulation, the “active ingredient” is the isopropylamine salt of glyphosate while the “other ingredients” include the surfactant polyethoxylated tallow amine (POEA) and other undisclosed chemicals (1,3).

Glyphosate is an amphoteric chemical that is often derived from the amino acid glycine (4,5). It is a weak organic acid with a water solubility of 12g/L at 25°C (6). Glyphosate and its salts are non-volatile, do not undergo photochemical degradation or hydrolysis at pH values of 3, 6, or 9 between 5°C and 35°C, and are detectable in soil after application for 1-151 days depending on soil types and environmental conditions (7–9). Although it is often reported as a single chemical, to increase its water solubility, it is produced for commercial use in a variety of salt forms including sodium, potassium, ammonium, isopropylammonium, and trimesium salts (10,11).

Glyphosate’s herbicidal action has been attributed to inhibiting the synthesis of aromatic amino acids through the shikemic acid pathway (12,13). This pathway is necessary for viability in plants, bacteria, fungi, and archaea. A common rationale for glyphosate’s safety was that it would be non-toxic to animals since the shikimic acid pathway is not found in animals (12,13). When it was first released for use in agriculture in the early to mid-1970s, glyphosate was hailed as “a once in a century herbicide” with the promise of decreasing the world’s total use of herbicides (1,10,14). By the end of the 1970s, glyphosate became the most commonly used herbicide in the world.

A number of developments helped glyphosate expand its dominance as the world’s most common herbicide, including: 1) the use of glyphosate-based herbicides (GBHs) in pre-emergent weed control in conjunction with no-till agricultural practices (2,15); 2) the development of genetically modified crops, such as corn and soybeans, that were resistant to Monsanto’s GBH Roundup^®^ and first marketed as “Roundup Ready^®^” in 1996 (2); 3) the expiration of the Monsanto patent and the subsequent production by other large agribusinesses such as *Dow AgroSciences*^®^ and *Syngenta*^®^ (3); and 4) the use of GBHs as desiccants to speed crop harvesting (2,16). An analysis by Benbrook in 2016 found that in the 40 years between 1974 and 2014, about two-thirds of all glyphosate applied in the US had been used in the last 10-year period (2).

The extensive amount of GBHs used in an expanding variety of agricultural practices when growing crops both resistant and not resistant to glyphosate has led to residues of glyphosate detected in human foods, beverages, cotton fabrics, and bandages as well as in human urine and breast milk (17–24). Studies performed by non-agricultural industry parties questioned the veracity of the safety conclusions of both governmental and agricultural industry groups. As examples, although the US EPA states that “glyphosate is no more than slightly toxic to birds and is practically nontoxic to fish, aquatic invertebrates, and honeybees”, studies have shown a higher toxicity towards amphibians and aquatic invertebrates (25–27). Glyphosate has been shown to change the behavior and reproduction of earthworms (28) and negatively affect the growth of algae and bacteria in aquatic systems (29). More recently, glyphosate has been shown to affect the gut microbiota of honeybees and is suggested as a possible contributing factor in colony collapse disorder (30,31). Glyphosate has been implicated in a variety of toxicity risks in vertebrate animals, including cell signal disruption in rats (32), endocrine disruption in human cell lines (33), and steroidogenesis disruption in the mouse MA-10 Leydig tumor cell line (34).

In addition to glyphosate’s direct toxicity, the unlisted ingredients in GBHs (frequently labeled “other ingredients”) have been shown to pose a risk to aquatic organisms (29,35). The EPA policy of not including safety testing for unlisted ingredients in pesticides has been called into question by studies of pesticide toxicity with and without the active ingredient (11,36,37). As reported in a publication on the toxicity of pesticides’ unlisted ingredients in *Scientific Reports*, “These adjuvants are generally considered by the EPA to be biologically inert; and therefore, their use is not monitored at the federal level and they are exempt from residue tolerance for food use” (38).

The unlisted ingredients in GBHs are trade secrets. The most prominent known component of the unlisted substances in Roundup^®^ is POEA, which is a surfactant that improves the penetration of glyphosate across the epidermis into plant tissues (39). POEA and other surfactants are not typically added to commercial preparations sold for use in aquatic environments, such as *Dow Agrosciences’* Rodeo^®^ aquatic herbicide (Rodeo), since surfactants are known to have harmful effects to aquatic organisms (40–42). Correspondence with Dow technical services confirmed that there are no surfactants present in Rodeo.

Since glyphosate is never applied as the sole ingredient in an herbicide, one must also consider the “other ingredients” to properly assess the toxicity of the full GBH product (10). To understand the safety, health, and environmental risks of GBHs, all their ingredients should be assessed, as well as the active ingredient in all its various chemical formulations.

A review of research regarding the effects of GBHs and their undisclosed ingredients on *D. magna* by Chura et.al. revealed a wide variety of approaches for toxicity with correspondingly varying conclusions (10). The effects have been monitored on development, egg production, survival rate, and heart rate (10) with a variety of culture waters, times monitored for survival, and sources of active and undisclosed ingredients. The culture water varies widely and includes synthetic water (43), Aachener Daphnien Medium (27), Elendt-M7 medium (40,41,44), and “moderately hard synthetic freshwater” (42). Survival assessments with EC50 or LC50 to the GBHs or glyphosate are typically at 24 or 48 hours with 48 appearing to be the most common (27,42,45). Tests regarding the toxicity of the GBHs’ surfactants or residues have generally used an EC50 or LC50 at 48 hours (40,42,45–47). The surfactants tested vary from those supplied by Monsanto (45) to general surfactants (40) and the known surfactant in Roundup^®^, POEA (42,48).

In this paper, we attempt to separate the effects of the active and undisclosed ingredients that may be involved in GBH toxicity by comparing the full GBH product to the known ingredients alone. We examined the effects on heart rate and survival over more discrete and shorter time periods than previously used. Our approach sought to mimic the natural environment and reduce stress during testing. Here, we report the comparative effects of “*Roundup Ready-to-Use Weed and Grass Killer*^®^”(Roundup-WGK), Rodeo, glyphosate, and POEA on the heart rates and survival rates of *Daphnia magna*. Our research seeks to determine whether Roundup-WGK and Rodeo, including their unlisted ingredients as well as their known components, glyphosate and POEA, have any deleterious effects on *D. magna*. *D. magna* is a fresh water aquatic invertebrate and a well-established model organism for toxicological studies (49,50). Because it is a filter feeder, it is rapidly responsive to suspended or dissolved substances, allowing for simple and efficient toxicological testing of chemicals (49,50). *D. magna* is also transparent and its heart rate can be directly observed with a stereo microscope (35). We hypothesized that the herbicide commercial formulations would be the most toxic and would affect both the heart rates and survival more than the individual components would by themselves.

## 2 Materials and methods

### 2.1 Materials

Adult *D. magna* were originally purchased from *Carolina Biological Supply* and grown in our lab at 19-22°C in two 10-gallon glass tanks filled with non-chlorinated artesian well water and fitted with aerators. Approximately 1/3^rd^ of the water was removed every 10-14 days and replaced with fresh non-chlorinated artesian well water previously allowed to equilibrate to room temperature. *D. magna* were maintained on a diet of spirulina (*Pure Planet*) and *Nannochloropsis (sp.)* unicellular green algae *(Carolina Biological Supply* Item #142337). A Wolfe^®^ stereomicroscope from *Carolina Biological Supply* was fitted with a custom camera adapter to capture videos of the *D. magna* at a magnification of 30x using an iPhone^®^ 6 Plus (*Apple^®^*). The microscope was modified with a cooling fan and LED illumination to control for temperature variation. Because of *magna*’s sensitivity to temperature change, the temperature was maintained within one degree of 21°C during testing. A Fluke^®^ 62 Max+ infrared thermometer with a resolution of 0.1°C was used to ensure that temperature was monitored accurately.

Four substances were tested: *Monsanto*’s Roundup-WGK containing 2% glyphosate isopropylamine salt and 98% unlisted ingredients including POEA, *Dow Agrosciences*’ Rodeo containing 53.8% glyphosate isopropylamine salt and 46.2% unlisted ingredients, pure glyphosate (*Sigma Aldrich*), and POEA (POE (15) tallow amine, *Chem Service, Inc*.).

All test substances were diluted with water from the *D. magna* culture tanks to create stock solutions. The choice of using tank water was based on our objective to minimize stress on the *D. magna* by avoiding a sudden change in their water. Most ready-to-use GBHs, including Roundup-WGK, contain 2% glyphosate salt and approximately 1% POEA (39). Five stock solutions were created that contained the percentages of glyphosate (2%) or POEA (1%) found in ready-to-use products. Since Rodeo’s recommended dilution results in a higher concentration of glyphosate, a sixth stock solution was created for Rodeo according to the manufacturer’s recommended dilution. The six stock solutions are:

1. **Roundup-WGK**: The original undiluted Roundup-WGK herbicide stock contained 2% glyphosate salt, an estimated 1% POEA, and 97% unlisted ingredients.
2. **Rodeo-Recommended**: The Rodeo stock contained 5.38% glyphosate, no surfactant or POEA, and 4.62% unlisted ingredients and was diluted from *Dow Agrosciences*’ *Rodeo* aquatic herbicide based on the manufacturer recommendation.
3. **Rodeo-2%**: The Rodeo-2% stock contained 2% glyphosate salt, no surfactant or POEA, and 1.7% unlisted ingredients and was diluted from *Dow Agrosciences*’ *Rodeo* aquatic herbicide. This solution was designed to match the glyphosate content in Roundup-WGK solution.
4. **Glyphosate**: The glyphosate stock contained 2% pure glyphosate.
5. **POEA**: The POEA stock contained 1% POEA.
6. **Mock-GBH**: The Mock-GBH contained 2% pure glyphosate and 1% POEA.

All stock solutions were diluted with water from the *D. magna* culture tanks to achieve concentrations ranging from 0.1% to 100% of the stock solution. A special 200% test concentration was created for both Rodeo-Recommended and Rodeo-2%. For comparison to other research using concentrations of glyphosate and/or POEA, we have provided a table with the concentrations (in mg/L) of each known ingredient in our dilution series (Table 1). Percent solutions were used since we wished to compare the effects of unlisted ingredients that have undeclared concentrations.

**Table 1.**
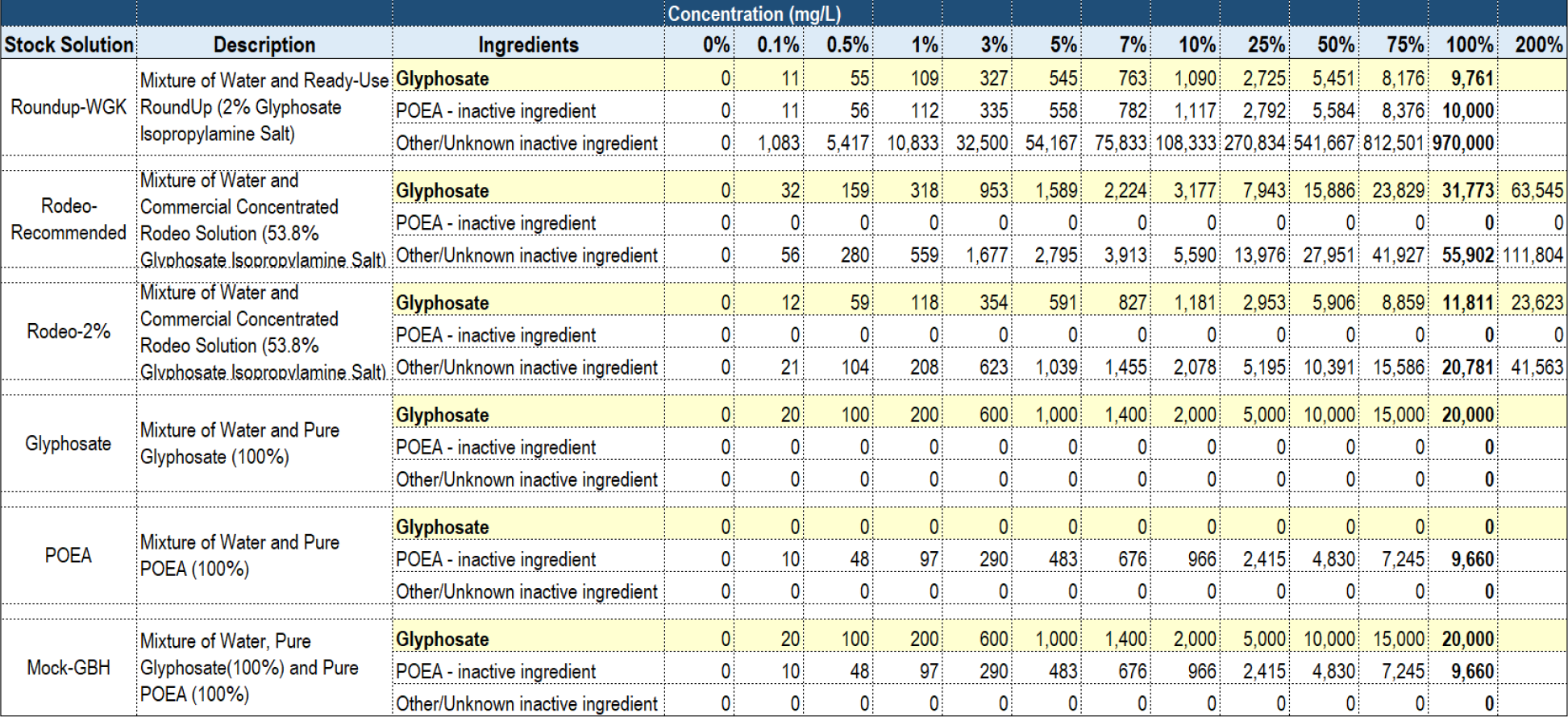
Concentration in mg/L of ingredients for each dilution of all stock solutions. Table 1 lists the mg/L of glyphosate, POEA, and “Other ingredients for the various percentage dilutions of the six stock solutions.

### 2.2 *Daphnia magna* heart rate analysis methods

Each experiment had a 0% control group containing water from *D. magna* culture tanks grown in our lab. Each *D. magna* was placed in an individual well of a 96-well microtiter plate filled with a given concentration of a test solution. Using our modified stereoscope, *D. magna* heart rates were quantified by videotaping each *D. magna* individually for 10 seconds on an iPhone 240 fps slow-motion setting and then manually counting their heart rates in the slow-motion video after the experiment was conducted. All heart rate measurements and analyses are derived from beats per minute (BPM). For all experiments, the heart rates of three *Daphnia magna* per condition were recorded.

For Roundup-WGK solutions, three experiments were conducted with narrowing concentrations of Roundup-WGK tested. The first Roundup-WGK experiment tested the concentrations 10%, 25%, 50%, 75%, and 100% and a video was captured of each *D. magna* every minute for 14 minutes. To further assess the procedure’s reliability and examine dose responses in more narrow concentration ranges, two more experiments were conducted. The second Roundup-WGK experiment tested the concentrations 1%, 3%, 5%, 7%, and 10%, and a video was captured of each *D. magna* every 3 minutes for 30 minutes. The third Roundup-WGK experiment tested the concentrations 0.1%, 0.5%, 1%, 5%, and 10%, and a video was captured of each *D. magna* every minute for 14 minutes.

For the test substances, Rodeo-Recommended, Rodeo-2%, glyphosate, POEA, and the Mock-GBH solutions, the concentrations 0.1%, 0.5%, 1%, 5%, 10%, 25%, 50%, 75%, and 100% were tested. An additional 200% concentration was tested for Rodeo-Recommended and Rodeo-2%. A video was captured of each *D. magna* every 5 minutes. For the concentrations 0.1%, 0.5%, 1% and 75% of Rodeo-Recommended and Rodeo-2% solutions, the observation duration was 35 minutes, whereas all others were 45 minutes. Table 2 displays the concentrations tested for each test substance.

**Table 2.**
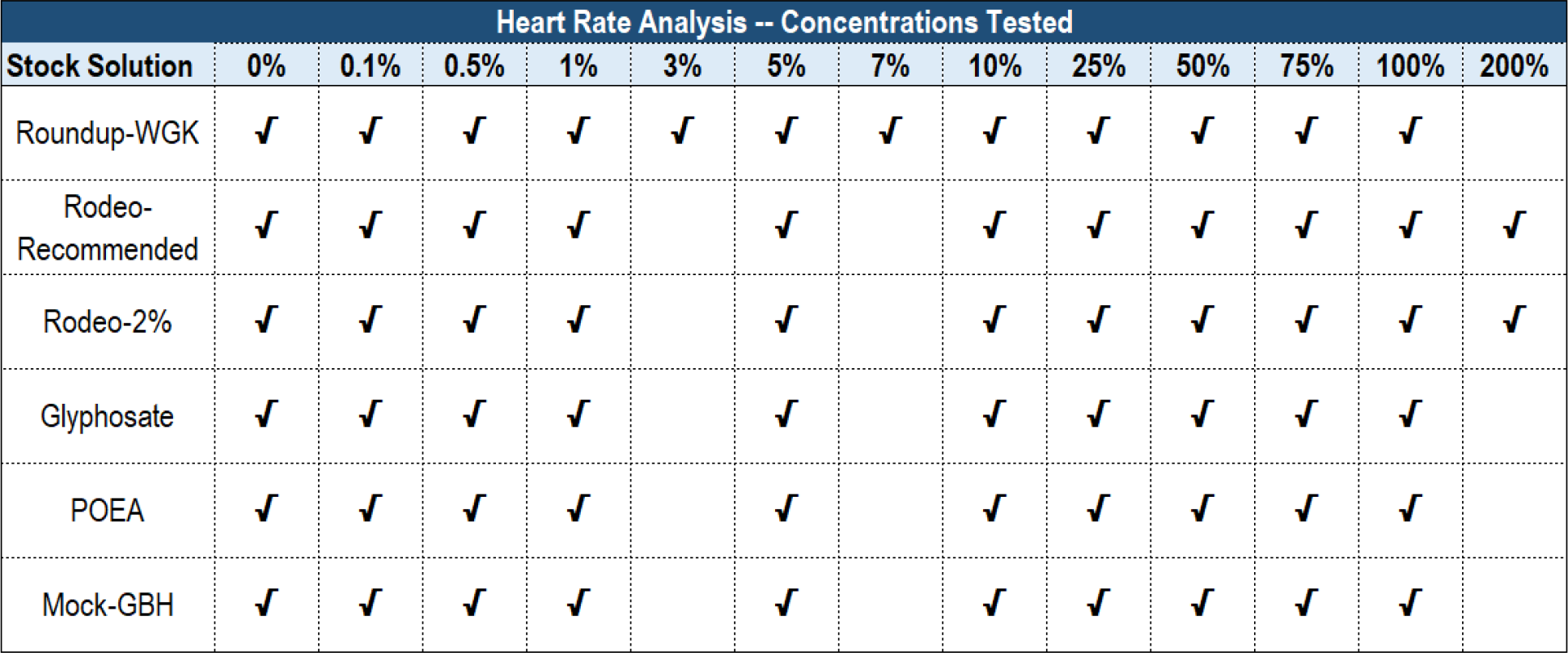
Concentrations tested for heart rate analyses. Table 2 shows the concentrations tested for Roundup-WGK, Rodeo-Recommended, Rodeo-2%, Glyphosate, POEA and Mock-GBH solutions. A check mark indicates that concentration was tested.

| Heart Rate Analysis -- Concentrations Tested |  |  |  |  |  |  |  |  |  |  |  |  |  |
| --- | --- | --- | --- | --- | --- | --- | --- | --- | --- | --- | --- | --- | --- |
| Stock Solution | 0% | 0.1% | 0.5% | 1% | 3% | 5% | 7% | 10% | 25% | 50% | 75% | 100% | 200% |
| Roundup-WGK | ✓ | ✓ | ✓ | ✓ | ✓ | ✓ | ✓ | ✓ | ✓ | ✓ | ✓ | ✓ |  |
| Rodeo-Recommended | ✓ | ✓ | ✓ | ✓ |  | ✓ |  | ✓ | ✓ | ✓ | ✓ | ✓ | ✓ |
| Rodeo-2% | ✓ | ✓ | ✓ | ✓ |  | ✓ |  | ✓ | ✓ | ✓ | ✓ | ✓ | ✓ |
| Glyphosate | ✓ | ✓ | ✓ | ✓ |  | ✓ |  | ✓ | ✓ | ✓ | ✓ | ✓ |  |
| POEA | ✓ | ✓ | ✓ | ✓ |  | ✓ |  | ✓ | ✓ | ✓ | ✓ | ✓ |  |
| Mock-GBH | ✓ | ✓ | ✓ | ✓ |  | ✓ |  | ✓ | ✓ | ✓ | ✓ | ✓ |  |

An expanded heart rate verification experiment was conducted to verify the results for Rodeo-Recommended, Rodeo-2%, and POEA. The concentrations 1%, 25%, and 100% were tested for each stock solution and ten *D. magna* were individually observed for each concentration. A video was captured of each *D. magna* every 15 minutes for 60 minutes.

Statistical analysis of the heart rate results was performed by applying a Kruskal-Wallis ANOVA test with Dunn’s Multiple Comparison post-test using GraphPad Prism version 8 for Mac (GraphPad Software, La Jolla, California, www.graphpad.com). Graphs were generated by plotting average heart rates with standard deviation using GraphPad Prism. Full statistical data is presented in Supporting Information (S1 Dataset), as is the raw data for the initial heart rate experiments (S2 Dataset) and heart rate verification experiments (S3 Dataset).

### 2.3 *Daphnia magna* survival rate analysis methods

To observe the effects of the test substances on *D. magna* survival, we used a larger containment vessel with 15-25mls of test solution in 25mm × 150mm glass tissue culture test tubes. Each experiment had a 0% control group. For each concentration, the time of death was recorded starting from when the first *D. magna* was introduced into a tube. Each tube was monitored continuously for eight hours. The ambient room temperature was kept between 19 and 22°C.

Two experiments were conducted on the Roundup-WGK solution. The concentration between 1% and 10% were tested. Each test concentration and control group contained twelve *D. magna* with 3 tubes per group and four *D. magna* per tube; 25mls of each concentration per test tube. Two experiments were conducted on Rodeo-Recommended and Rodeo-2% solutions. One experiment was conducted on glyphosate, POEA, and Mock-GBH solutions. For Rodeo-Recommended, concentrations between 1% and 200% were tested. For Rodeo-2%, concentrations between 1% and 75% were tested. For glyphosate, POEA and Mock-GBH, concentrations between 1% and 100% were tested. Each test concentration and control group contained 5-6 *D. magna* with 3 tubes per group and 1 to 2 *D. magna* per tube; 15mls of each concentration per test tube. Table 3 displays the concentrations tested for each test substance.

**Table 3.** Concentrations tested for survival rate analysis. Table 3 outlines the concentrations tested of Roundup-WGK, Rodeo-Recommended, Rodeo-2%, Glyphosate, POEA and Mock-GBH solutions for survival analysis. A check mark indicates that the concentration was tested.

| Median Time Until Death Analysis -- Concentrations Tested |  |  |  |  |  |  |  |  |  |  |  |  |  |  |
| --- | --- | --- | --- | --- | --- | --- | --- | --- | --- | --- | --- | --- | --- | --- |
| Stock Solution | 0% | 1% | 2% | 3% | 4% | 5% | 7.50% | 10% | 25% | 30% | 50% | 75% | 100% | 200% |
| Roundup-WGK (12D, 2017-#1) | ✓ | ✓ | ✓ | ✓ | ✓ |  |  |  |  |  |  |  |  |  |
| (12D, 2017-#3) | ✓ |  |  |  |  | ✓ | ✓ | ✓ |  |  |  |  |  |  |
| Rodeo-Recom. (5D, 2017-02-04) | ✓ | ✓ |  |  |  | ✓ |  | ✓ |  |  |  |  | ✓ | ✓ |
| (6D, 2017-12-20) | ✓ |  |  |  |  |  |  |  | ✓ |  | ✓ | ✓ |  |  |
| Rodeo-2% (5D, 2017-02-04) | ✓ | ✓ |  | ✓ |  | ✓ |  | ✓ |  | ✓ |  |  |  |  |
| (6D, 2017-12-20) | ✓ |  |  |  |  |  |  |  | ✓ |  | ✓ | ✓ |  |  |
| Glyphosate (6D, 2017-07-18) | ✓ | ✓ |  | ✓ |  |  |  | ✓ |  | ✓ |  |  | ✓ |  |
| POEA (6D, 2017-07-18) | ✓ | ✓ |  |  |  | ✓ |  | ✓ | ✓ |  | ✓ |  | ✓ |  |
| Mock-GBH (6D, 2017-07-18) | ✓ | ✓ |  |  |  | ✓ |  | ✓ | ✓ |  | ✓ | ✓ | ✓ |  |

Kaplan-Meier survival curves were generated in Microsoft Excel and median time until death was determined manually for all concentrations. The survival rate raw data is provided in the Supporting Information (S4 Dataset).

## 3 Results

### 3.1 Daphnia magna heart rate analysis

Heart rate experiments were performed with a 0% control. There is a wide range of resting heart rates for control groups, ranging 126-546 BPM at a temperature of 21°C. Supplemental Figure 1 (S1 Fig) shows the compiled average and median heart rates of all the control groups in our heart rate experiments with each data point representing 25 *D. magna* (S1 Fig). Unlike test groups, control heart rates remained steady without dramatic decreases or increases. The median and mean heart rates of *D. magna* in our experiments were 369 and 357 BPM respectively. The consolidated control heart rate raw data is provided in the Supporting Information (S5 Dataset).

**Figure 1.**
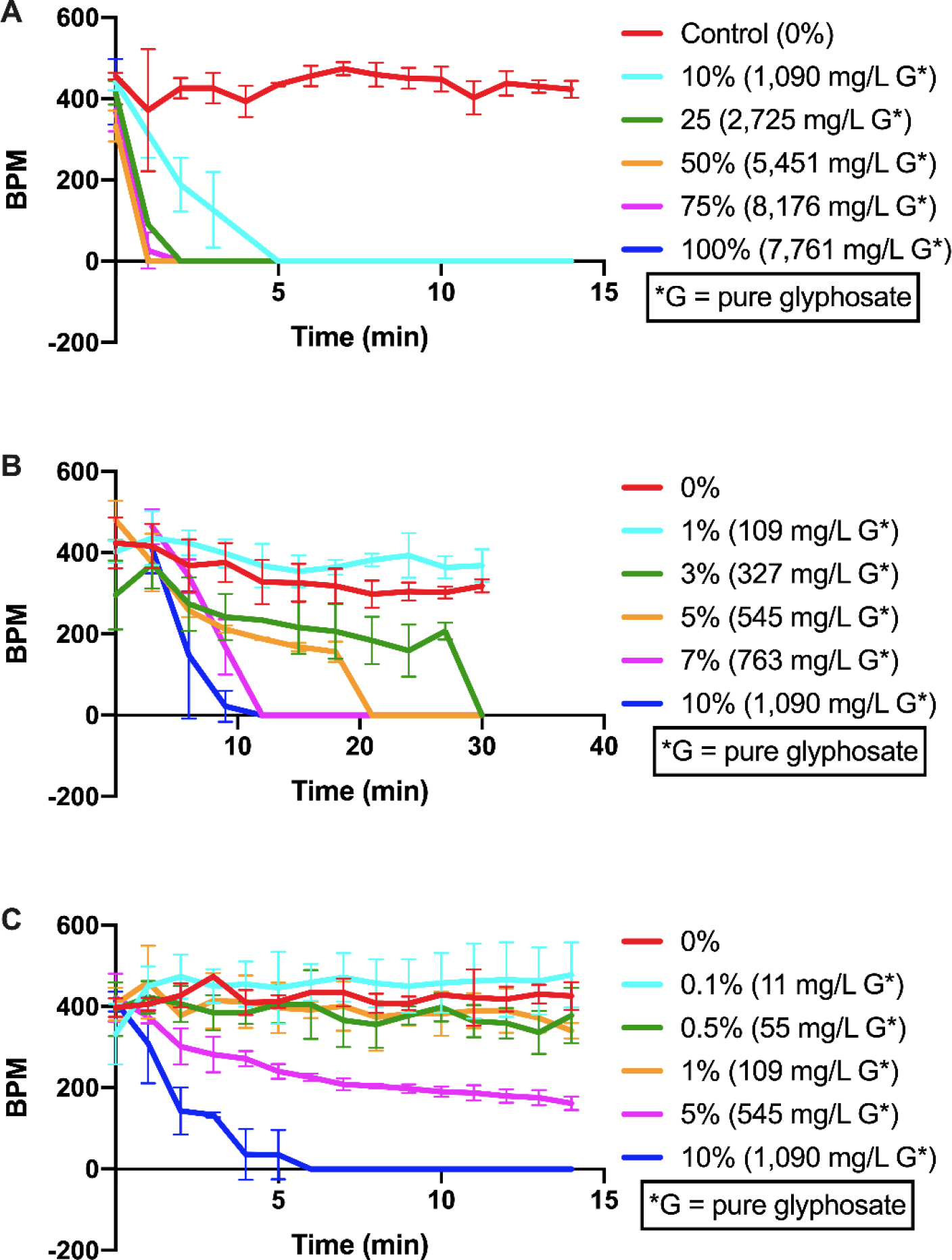
Effects of Roundup-WGK solutions on heart rate. Graphs represent the average BPM with standard deviation of the *D. magna* over exposure time in minutes for listed concentrations. Each data point represents 3 *D. magna*. The control (0% Roundup-WGK) is shown as a red line. **A)** The first experiment looked at a broad range of concentrations: 0%, 10%, 25%, 50%, 75%, and 100% Roundup-WGK (100% stock contains 2% glyphosate, approximately 1% POEA). Heart rates show a precipitous decline for all concentrations tested, following a dose-response pattern. All test concentrations yielded an average heart rate significantly below that of the control (p<0.001) **B)** The second experiment focused on the range of 0-10% with concentrations of 1%, 3%, 5%, 7%, and 10% Roundup-WGK. Heart rates showed a clear dose response above 1% Roundup-WGK. The *D. magna* in the 7% and 10% concentrations had average heart rates significantly below that of the control group (p<0.05) **C)** The last experiment tested lower concentrations of 0.1%, 0.5%, 1%, 5% and 10% Roundup-WGK. Heart rates showed a clear dose response above 1% Roundup-WGK, with 5% and 10% being significantly decreased compared to the control (p<0.05). *D. magna* in the 0.1%, 0.5%, and 1% solutions had heart rates within the range set by the controls.

#### 3.1.1 Roundup-WGK

We first performed a study examining the effects of successively more narrow concentration ranges of Roundup-WGK on heart rate. For the 10% to 100% concentrations, *D. magna* heart rates dropped to 0 BPM within 8 minutes (Fig 1A) with an average heart rate significantly lower (p<0.001) than the control. To further explore the effects of concentrations between 0% and 10%, we performed two more experiments. In the second experiment, *D. magna* heart rates dropped to 0 BPM in less than 30 minutes for the 3%, 5%, 7%, and 10% concentrations (Fig 1B). In the third experiment, with lower concentrations, *D. magna* heart rates again dropped to 0 BPM in less than 30 minutes for the 5% and 10% concentrations. For the 0.1%, 0.5%, and 1% concentrations, *D. magna* heart rates remained within the normal range set by the control (Fig 1C).

#### 3.1.2 Rodeo-Recommended and Rodeo-2%

The 5% to 25% solutions reduced their heart rates to approximately 75% of the control within 45 minutes (Fig 2A). The 75% to 100% concentrations of the Rodeo-Recommended stock solution reduced the *D. magna*’s heart rate by at least 50% of the control within 45 minutes (Figs 2A and 2C). The 200% solution reduced their heart rates to approximately 20% of the control (Fig 2B). The 0.1% and 0.5% concentrations remained within the normal range set by the control. No death was observed within the 45-minute observation period (Fig 2C).

**Fig 2.**
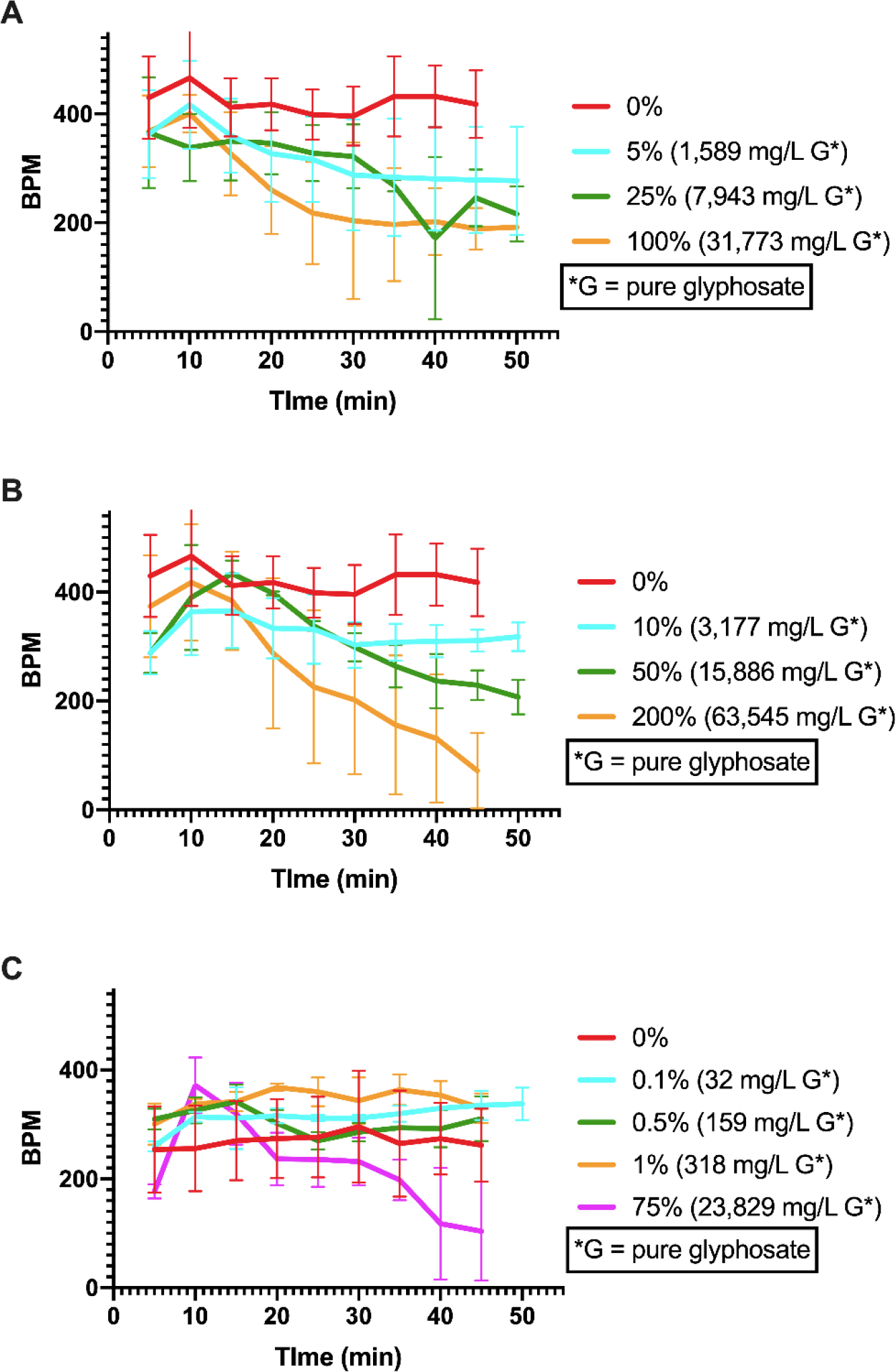
Effects of Rodeo-Recommended solutions on heart rate. This represents the average BPM with standard deviation of the *D. magna* over exposure time in minutes for listed concentrations of the Rodeo-Recommended stock (100% stock contains 5.38% glyphosate). The control (0%) is shown as a red line. Each data point represents 3 *D. magna*. **A)** Heart rates at the concentrations 0%, 5%, 25%, and 100%. **B)** Heart rates at the concentrations 0%, 10%, 50%, and 200%. Graphs **A** and **B** represent data from a single experiment, separated for easier viewing, and share the same control. **C)** Heart rates at the concentrations 0%, 0.1%, 0.5%, 1%, and 75%. There was extensive overlap for the lower concentrations. The higher concentrations of 75%, 100%, and 200% showed a clear decline by 30 minutes that continued out to 45 minutes. Average heart rates for all concentrations ≥ 1% Rodeo-Recommended showed a significant difference from control except for the 75% test group (p<0.05).

Heart rates approximated a general dose-response pattern with higher concentrations separating from the lower concentrations by the end of the 45-minute observation period. Extensive crossover among test groups is observed at earlier time points. Average heart rates in all concentrations ≥1% Rodeo-Recommended varied significantly from the control heart rates (p<0.05) except the 75% group, which varied widely around the control line, ending at less than 50% of the control (Figs 2A, 2B and 2C).

Heart rates ranged from 100% to 60% of the control for the 100% to 0.1% concentrations within 45 minutes, approximating a general dose-response pattern. No death was observed within 45 minutes (Figs 3A, 3B and 3C). For the Rodeo-2% stock, the 200% concentration reduced the heart rates to approximately 50% of the control within 45 minutes (Fig 3B).

**Fig 3.**
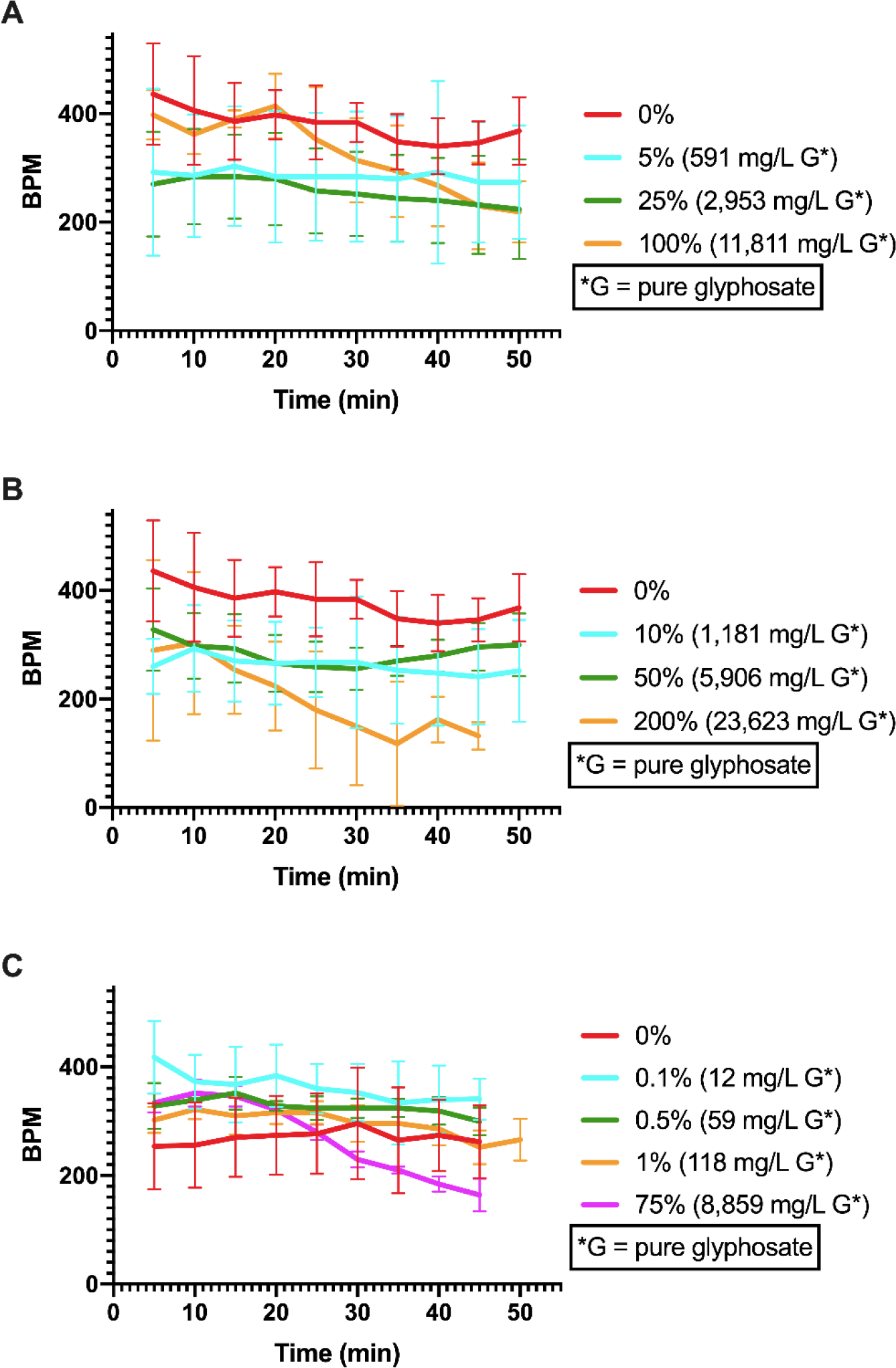
Effects of Rodeo-2% solutions on heart rate. This represents the average BPM with standard deviation of the *D. magna* over exposure time in minutes for listed concentrations of the Rodeo-2% stock (100% stock contains 2% glyphosate). The control (0%) is shown as a red line. Each data point represents 3 *D. magna*. **A)** Heart rates in the concentrations 0%, 5%, 25%, and 100%. **B)** shows the concentrations 0%, 10%, 50%, and 200%. Graphs **A** and **B** represent data from a single experiment, separated for easier viewing, and share the same control. Each graph contains a range of high and low concentrations for ease of comparison. **C)** Heart rates in the concentrations 0%, 0.1%, 0.5%, 1%, and 75%. With the exception of the 50%, there was a general trend for the higher concentrations of 25% to 200% to show decreasing heart rates in a dose-response pattern. The 0.1%, 0.5%, 10%, 25%, 50%, and 200% Rodeo-2% group heart rates varied significantly compared to the control (p<0.05).

With the exception of the 50%, there was a general trend for the higher concentrations of 25% to 200% to show decreasing heart rates in a dose-response pattern. Average heart rates in the 0.1%, 0.5%, 10%, 25%, 50%, and 200% Rodeo-2% groups were significantly different from control heart rates (p<0.05) (Figs 3A, 3B and 3C).

#### 3.1.3 Glyphosate

For most concentrations >1% glyphosate, heart rates of *D. magna* varied from the control group. However, the heart rates remained steady and did not show a dose-response pattern relative to the control. Average heart rates in 5%, 10%, 25%, 50%, and 75% glyphosate groups were significantly different from control heart rates (p<0.05) (Figs 4A and 4B).

**Fig 4.**
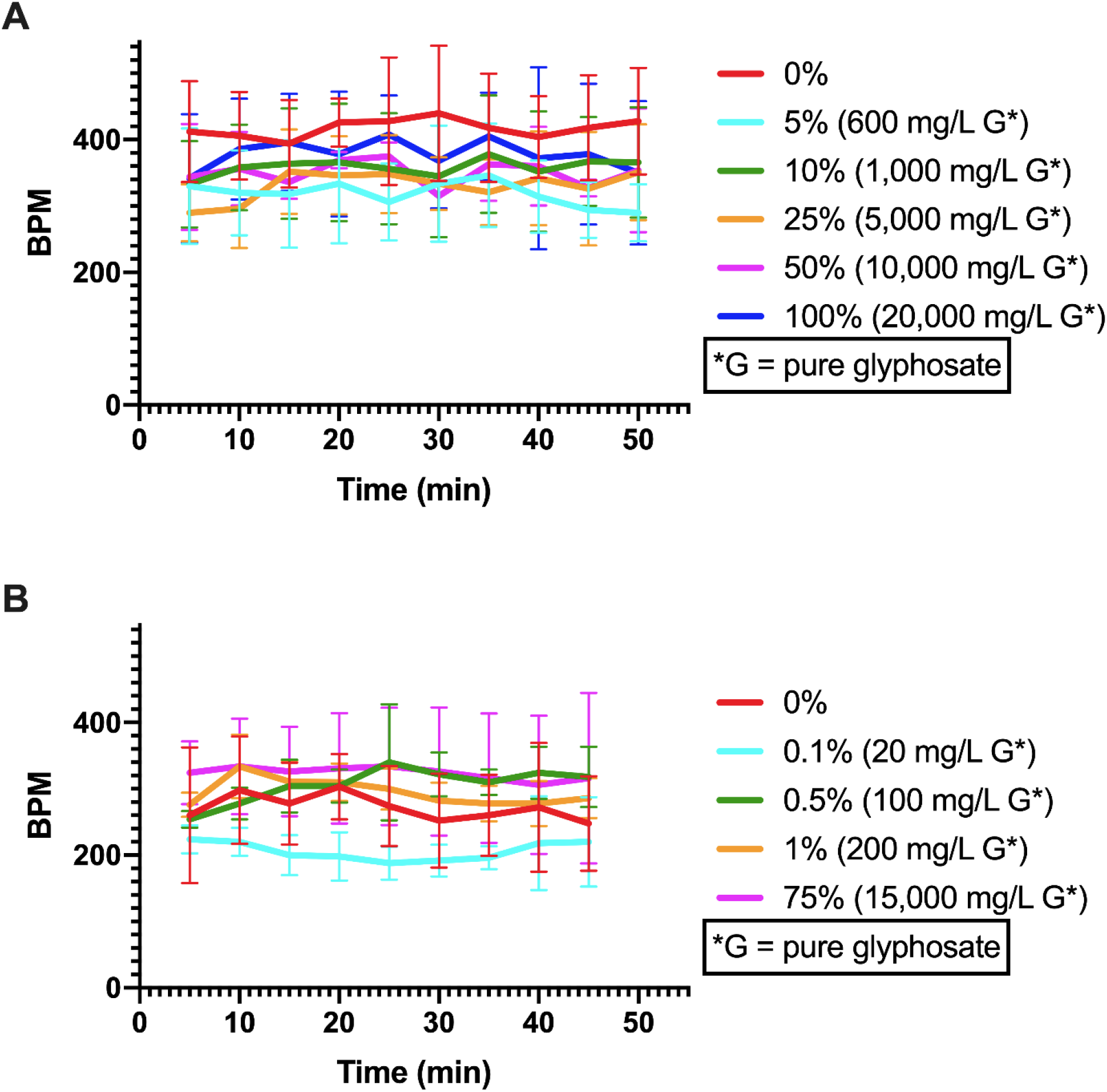
Effects of glyphosate solutions on heart rate. This represents the average BPM with standard deviation of the *D. magna* over exposure time in minutes for listed concentrations of glyphosate (100% stock contains 2% glyphosate). The control (0%) is shown as a red line. Each data point represents 3 *D. magna*. Concentrations are divided into 2 graphs for clarity and to match the concentration groupings of the previous experiments. All concentrations presented here share the same control. **A)** Heart rates in the concentrations 0%, 5%, 10%, 25%, 50%, and 100%. **B)** Heart rates in the concentrations 0%, 0.1%, 0.5%, 1%, and 75%. For the tested concentrations, the heart rates remained steady and did not show a dose-response pattern. The heart rates of *D. magna* in the 5%, 10%, 25%, 50%, and 75% glyphosate groups were significantly different from those of the controls (p<0.05).

#### 3.1.4 POEA

In the POEA test groups, the *D. magna* heart rates ranged from approximately 15% to 100% of the control group, following no clear dose response (Figs 5A and 5B). The 5% concentration showed steep jump in heart rate (Fig 5A). Average heart rates in all solutions ≥1% except for the 25% POEA solution were significantly lower than control heart rates (p<0.05). Though there is no clear dose-response, the *D. magna* in the three highest concentrations of 50%, 75% and 100% had distinctly lower heart rates (p<0.0001) at less than 200 BPM (Figs 5A and 5B).

**Fig 5.**
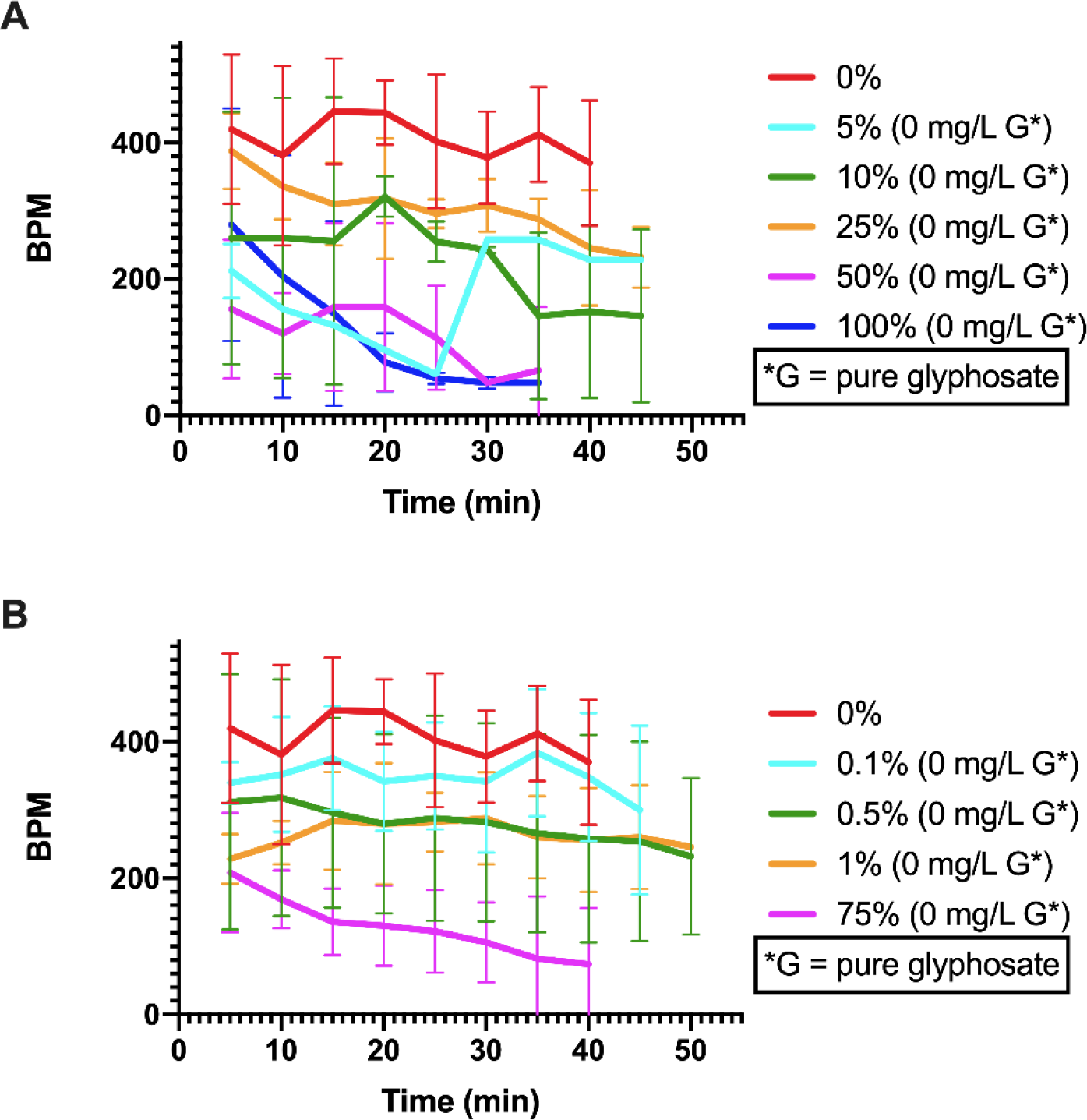
Effects of POEA solutions on heart rate. This represents the average BPM with standard deviation of the *D. magna* over exposure time in minutes for listed concentrations of POEA (100% stock contains 1% POEA and no glyphosate). The control (0%) is shown as a red line. Each data point represents 3 *D. magna*. **A)** Heart rates in the concentrations 0%, 5%, 10%, 25%, 50%, and 100%. **B)** Heart rates in the concentrations 0%, 0.1%, 0.5%, 1%, and 75%. Concentrations are divided into 2 graphs for clarity and to match the concentration groupings of the previous experiments. All concentrations presented here share the same control. Although all test concentrations had heart rates lower than controls, there was no clear dose-response pattern. However, the 3 highest concentrations of POEA prompted the most drastic decreases in heart rate (p<0.0001). All concentrations ≥1% of the POEA stock solution, except for the 25% solution, produced heart rates significantly lower than the control group (p<0.05). Missing data points were the result of unreadable videos. This is generally revealed in Figs 5A and 5B where lines do not continue out to 45 minutes. For all missing data points, see highlighted data fields in S2 Dataset.

#### 3.1.5 Mock-GBH (Glyphosate + POEA)

Heart rates following exposure to Mock-GBH concentrations decreased over time compared to the control except for in the 0.5% concentration. The heart rates did not follow a dose-response pattern and ranged from 150% to 25% of the control. Average heart rates in the 10%, 25%, and 75% Mock-GBH groups were significantly lower than control heart rates (p<0.05) (Figs 6A and 6B).

**Fig 6.**
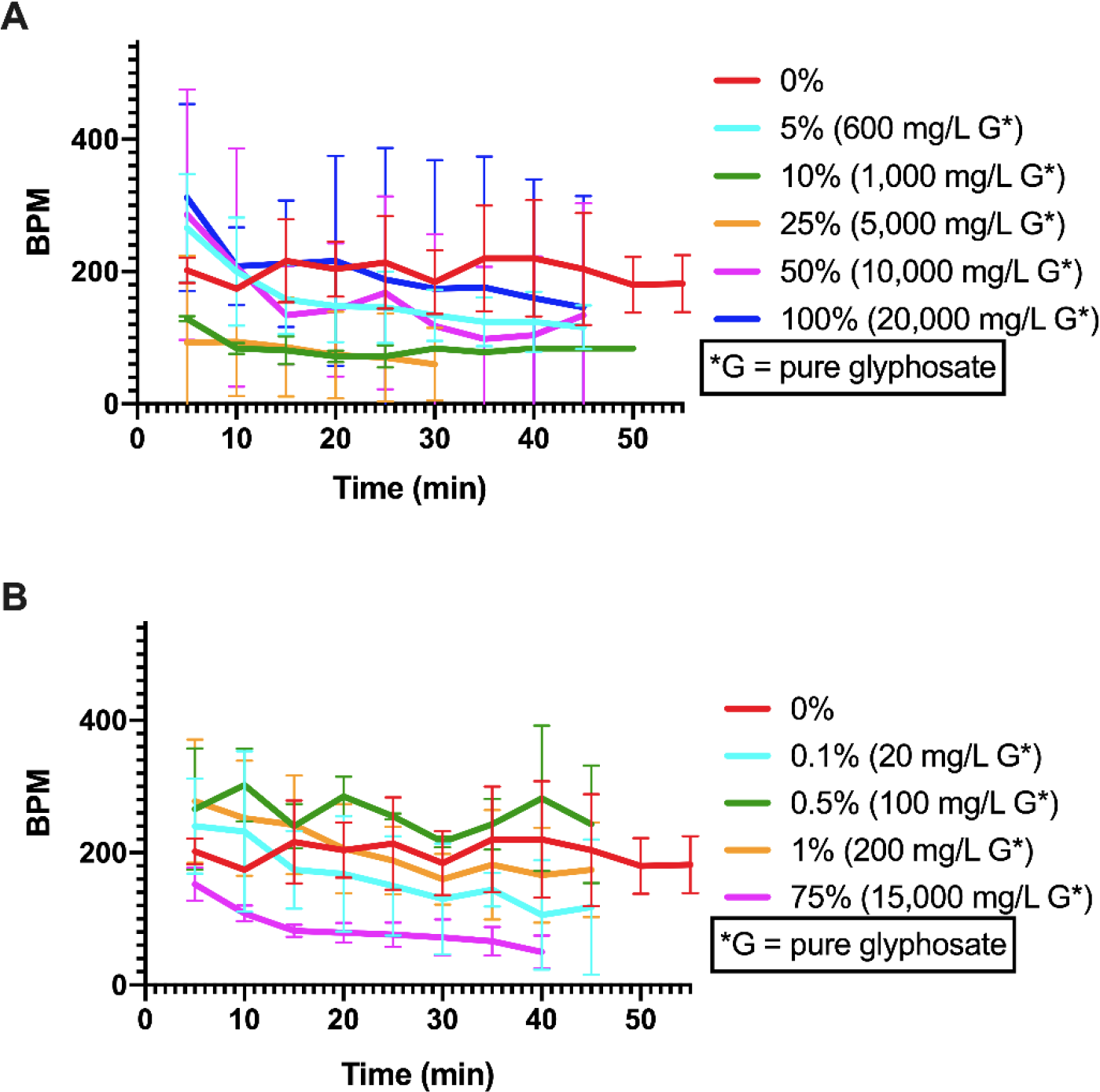
Effect of Mock-GBH (glyphosate + POEA) solutions on heart rate. This represents the average BPM with standard deviation of the *D. magna* over exposure time in minutes for listed concentrations of the Mock-GBH (100% stock contains 1% POEA and 2% glyphosate). The control (0%) is shown as a red line. Each data point represents 3 *D. magna*. **A)** Heart rates for the concentrations 0%, 5%, 10%, 25%, 50%, and 100%. **B)** Heart rates for the concentrations 0%, 0.1%, 0.5%, 1%, and 75%. Concentrations are divided into 2 graphs for clarity and to match the concentration groupings of the previous experiments. All concentrations presented here share the same control. The control in this experiment had heart rates approximately 50% slower than the heart rates of controls in other experiments. Heart rates did not follow a dose-response pattern, although *D. magna* in the 10%, 25%, and 75% Mock-GBH concentrations had an average heart rate significantly lower than that of the control. Missing data points were the result of unreadable videos. This is generally revealed in Figs 6A and 6B where lines do not continue out to 45 minutes. For all missing data points, see highlighted data fields in S2 Dataset.

All *D. magna* test and control groups displayed lower heart rates than in all other experiments of this investigation. This is revealed by all concentrations settling at under 200 BPM, with exception of the 0.5% concentration (Figs 6A and 6B).

#### 3.1.6 Verification experiments for Rodeo-Recommended, Rodeo-2%, and POEA

The results for Rodeo-Recommended, Rodeo-2%, and POEA did not follow clear dose-response patterns (Figs 2A,B,C, 3A,B,C and 5A,B). This led us to perform verification experiments for the *D. magna* heart rates in Rodeo-Recommended, Rodeo-2%, and POEA solutions. The concentrations 1%, 25%, and 100% were chosen and a larger test group of 10 *D. magna* was used. *D. magna* heart rates were captured every 15 minutes for one hour.

The 1% and 25% concentrations of both the Rodeo-Recommended and Rodeo-2% stocks stayed within the normal range set by the control (S2 Figs A and B). Although the 100% concentrations of both Rodeo-Recommended and Rodeo-2% stock solutions caused the *D. magna* heart rates to decrease to less than 50% of the control by 60 minutes (S2 Figs A and B), only the Rodeo-2% results at 100% concentration were statistically significant from the control (p<0.05) (S2 Fig B).

The heart rates for the 1% concentration of the POEA stock remained within the normal range set by the control (S2 Fig C). The 25% and the 100% concentrations had lower heart rates at less than 50% of the control for the entire 60-minute observation period (S2 Fig C). Unlike the 100% concentrations of Rodeo-Recommended and Rodeo-2% (S2 Figs A and B), the POEA concentrations did not cause any sharp changes in heart rates over time (S2 Fig C). POEA heart rates remained steady and level, but with significantly (p<0.05) reduced BPM for the 25% and 100% concentrations. (S2 Fig. C).

### 3.2 Daphnia magna survival rate analysis

All survival rate experiments were performed with a control concentration of 0%. Controls had 100% survival with no control *D. magna* dying within the observation period of 8 hours. Survival results are summarized in Table 4 using median time until death. Kaplan-Meier plots of survival over time are supplied in the Supporting Information (S3–8 Figs).

**Table 4.** Median time until death of *D. magna* exposed to Roundup, Rodeo, glyphosate, POEA, and Mock-GBH solutions. Median time until death indicates the time at which half of the population died. “NA” indicates that fewer than half of the population died by the end of the experiment. A gray box indicates that the concentration was not tested. N=12 *D. magna* per concentration for Roundup-WGK; N=5 *D. magna* per concentration for 1%-10%, 100% and 200% of Rodeo-Recommended stock solution; and 1%-10% and 30% of Rodeo-2% stock solution; and N=6 *D. magna* per concentration for the rest of the stock solutions. For POEA and Mock GBH, all concentrations were tested on the same day. For Roundup, 1-4% were tested on one day and 5-10% tested on another. For Rodeo-Recommended and Rodeo-2%, the 25%, 50%, and 75% concentrations were tested on a separate day from the rest. All control *D. magna* survived for the entirety of the observation period of 8hrs. Kaplan-Meier plots of survival over time are supplied in the Supporting Information (S3–8 Figs).

| Stock Solution | Median Time Until Death (Hour : Minute : Second) |  |  |  |  |  |  |  |  |  |  |  |  |  |
| --- | --- | --- | --- | --- | --- | --- | --- | --- | --- | --- | --- | --- | --- | --- |
|  | 0% | 1% | 2% | 3% | 4% | 5% | 7.50% | 10% | 25% | 30% | 50% | 75% | 100% | 200% |
| Roundup-WGK (12D, 2017-#1) | NA | NA | 0:55:13 | 0:24:21 | 0:17:53 |  |  |  |  |  |  |  |  |  |
| (12D, 2017-#3) | NA |  |  |  |  | 0:08:50 | 0:03:40 | 0:02:59 |  |  |  |  |  |  |
| Rodeo-Rec. (5D, 2017-02-04) | NA | NA |  |  |  | NA |  | NA |  |  |  |  | 0:37:00 | 0:20:00 |
| (6D, 2017-12-20) | NA |  |  |  |  |  |  |  | 3:14:00 |  | 2:06:00 | 1:08:00 |  |  |
| Rodeo-2% (5D, 2017-02-04) | NA | NA |  | NA |  | NA |  | NA |  | NA |  |  |  |  |
| (6D, 2017-12-20) | NA |  |  |  |  |  |  |  | 6:32:00 |  | 2:40:00 | 2:19:00 |  |  |
| Glyphosate (6D, 2017-07-18) | NA | NA |  | NA |  |  |  | NA |  | NA |  |  | NA |  |
| POEA (6D, 2017-07-18) | NA | 5:18:00 |  |  |  | NA |  | NA | 3:49:00 |  | 2:52:00 |  | 2:16:00 |  |
| Mock-GBH (6D, 2017-07-18) | NA | NA |  |  |  | NA |  | 2:30:00 | NA |  | 4:22:00 | NA | 7:23:00 |  |

#### 3.2.1 Roundup-WGK

The 7.5% and 10% Roundup-WGK groups all *D. magna* died within 10 minutes (median time until death of 3min 40sec and 2min 59sec respectively) (Table 4, S3 Fig). All of the 5% group died within 12 minutes (median time until death 8min 50sec). The 4% group all died within 25min (median time until death 17min 53sec). The 3% group all died by 40 minutes (median time until death 24min 21sec) and the 2% group all died within 110 minutes (median time until death 55min 31sec) (Table 4, S3 Fig). In the 1% group, one *D. magna* died within 5 minutes, the remaining *D. magna* did not die within the 8-hour observation period.

#### 3.2.2 Rodeo-Recommended and Rodeo-2%

All *D. magna* died within 2 hours in the 100% and 200% concentrations of the Rodeo-Recommended stock (median time until death 37min and 20min, respectively) (Table 4, S4 Fig). The 75% group died within 3 hours (median time until death 1hr 8min), the 50% group died within 4 hours (median time until death 2hr 6min), and the 25% group died within 6 hours (median time until death 3hr 14min). The 1%, 5%, and 10% groups did not die within the 8-hour observation period (Table 4, S4 Fig.). Starting at 25%, death rates show a precipitous decline with a clear dose-response pattern (S4 Fig).

For the Rodeo-2% stock, the 75% and the 50% groups all *D. magna* died within 4 and 5 hours, respectively (median time until death 2hr 19min and 2hr 40min, respectively) (Table 4, S5 Fig). Half of the 25% group died within 8 hrs. The remaining groups of 10% down to 1% concentrations showed no deaths within the 8-hour period (Table 4, S5 Fig). Survival rates followed a dose-response pattern starting at the 25% concentration with the two highest concentrations showing a precipitous decline within 5 hours (S5 Fig).

#### 3.2.3 Glyphosate

No deaths were observed in any of the glyphosate test groups (Table 4, S6 Fig).

#### 3.2.4 POEA

The *D. magna* in the three highest concentrations of 100%, 50%, and 25% declined to 50% of control survival within the first 4 hours (median time until death 2hr 16min, 2hr 52 min, and 3hr 49 min, respectively). All of the *D. magna* died within 7 hours in the 100% concentration group. Half of the 1% group died within 6 hours (median time until death 5hr 18min), but less than half of the 5% and 10% groups died within the 8-hour observation period (Table 4, S7 Fig). Survival rates follow a dose-response pattern for the three highest concentrations dropping to 50% survival within the first 4 hours. The 1%, 5%, and 10% concentrations did not follow a dose-response pattern (S7 Fig).

#### 3.2.5 Mock-GBH (POEA + Glyphosate)

The Mock-GBH groups did not follow a dose-response pattern (Table 4, S8 Fig). All *D. magna* in the 50% group died within 8 hours (median time until death 4hr 22min). Approximately 67% of the *D. magna* died in the 10% group within 8 hours (median time until death 2hr 30min) and about 50% died within 8 hours in the 100% group (median 3 time until death 7hr 23min). For all other groups of 1%, 5%, 25%, and 75%, about 33% died within 8 hours (Table 4, S8 Fig). Survival rates do not follow a dose-response pattern (S8 Fig).

## 4 Discussion

An important consideration in this study is the sensitivity of *D. magna* species’ heart rates to variations in water temperature. For example, the heart rate of *D. magna pulex* increases by about 24 BPM per 1°C (51). Because water temperature was maintained at 21°C during all experiments, temperature had minimal effects on our results. The variation in our control heart rates is not uncommon and aligns with published data of normal adult *D. magna* heart rates at about 21°C (51,52). At this temperature, various publications have stated that *D. magna* heart rate ranges from approximately 180 to 350 BPM (50,52). Individual heart rates for *D. magna* species are also highly variable among individuals and across conditions (51). We are unsure why the control and experimental heart rates in the Mock-GBH experiment were unusually low compared to our other experiments presented in this paper.

As summarized in Table 5, glyphosate alone had the least effect on *D. magna* heart rates out of all test substances and showed no effect on survival rates (Fig 4, Table 4, and S6 Fig). Although POEA lowered heart and survival rates (Fig 4, Table 4, and S7 Fig), the lack of a clear dose response makes definitive conclusions difficult. POEA and glyphosate together in the Mock-GBH had approximately the same effect on heart rate and survival rate as POEA alone (Figs 5–6, Table 4, and S7–8 Figs), reinforcing the conclusion that glyphosate has a marginal effect on *D. magna* physiology at the concentrations used in this study. For the glyphosate, POEA, and Mock-GBH solutions, 100% concentrations are closest to the concentrations of these chemicals (in mg/L) in ready-to-use GBHs. Considering the unlisted ingredient concentration(s) are unknown, we diluted the full products for dose response testing using percent rather than diluting to a specific concentration in weight/volume of glyphosate or POEA. These results emphasize the probable deleterious effects of unlisted ingredients in herbicides.

**Table 5.** Summary of testing results. A qualitative summary of the effects of Roundup-WGK, Rodeo-Recommended, Rodeo-2%, Glyphosate, POEA, and Mock-GBH solutions on *D. magna* survival rates and heart rates. The x, xx, xxx, xxxx approximate increasing severity of response seen for those stock solutions.

| Stock Solution | Description | Ingredients | Percentage (V/V) | Concentration (g/L) | Impacts |  |
| --- | --- | --- | --- | --- | --- | --- |
|  |  |  |  |  | Survival Rate Reduction | Heart Rate Reduction |
| Roundup-WGK | Mixture of Water and Ready-Use RoundUp (2% Glyphosate Isopropylamine Salt) | Glyphosate Isopropylamine Salt | 2.00% | 20.0 | xxxx | xxxx |
|  |  | **Glyphosate** | 0.98% | 9.8 |  |  |
|  |  | POEA - inactive ingredient | 1.00% | 10.0 |  |  |
|  |  | Other/Unknown inactive ingredient | 97.00% | 970.0 |  |  |
| Rodeo-Recommended | Mixture of Water and Commercial Concentrated Rodeo Solution (53.8% Glyphosate Isopropylamine Salt) | Glyphosate Isopropylamine Salt | 5.38% | 65.1 | xxx | xxx |
|  |  | **Glyphosate** | 2.63% | 31.8 |  |  |
|  |  | POEA - inactive ingredient | 0.00% | 0.0 |  |  |
|  |  | Other/Unknown inactive ingredient | 4.62% | 55.9 |  |  |
| Rodeo-2% | Mixture of Water and Commercial Concentrated Rodeo Solution (53.8% Glyphosate Isopropylamine Salt) | Glyphosate Isopropylamine Salt | 2.00% | 24.2 | xx | xx |
|  |  | **Glyphosate** | 0.98% | 11.8 |  |  |
|  |  | POEA - inactive ingredient | 0.00% | 0.0 |  |  |
|  |  | Other/Unknown inactive ingredient | 1.7175% | 20.8 |  |  |
| Glyphosate | Mixture of Water and Pure Glyphosate (100%) | **Glyphosate** | 2.00% | 20.0 | None | None |
|  |  | POEA - inactive ingredient | 0.00% | 0.0 |  |  |
|  |  | Other/Unknown inactive ingredient | 0.00% | 0.0 |  |  |
| POEA | Mixture of Water and Pure POEA (100%) | **Glyphosate** | 0.00% | 0.0 | x | x |
|  |  | POEA - inactive ingredient | 1.00% | 9.7 |  |  |
|  |  | Other/Unknown inactive ingredient | 0.00% | 0.0 |  |  |
| Mock-GBH | Mixture of Water, Pure Glyphosate(100%), and Pure POEA (100%) | **Glyphosate** | 2.00% | 20.0 | x | x |
|  |  | POEA - inactive ingredient | 1.00% | 9.7 |  |  |
|  |  | Other/Unknown inactive ingredient | 0.00% | 0.0 |  |  |

As an internal verification of our experiments, Rodeo-2% containing 2% glyphosate has approximately half the concentration of glyphosate as is present in Rodeo-Recommended containing 5.38% glyphosate. The unlisted ingredients in Rodeo are also approximately halved. The decreased death rates for Rodeo-2% compared to Rodeo-Recommended support this pattern (Table 4, S4–5 Figs). The median time until death is approximately half in the Rodeo-2% compared to the Rodeo-Recommended (Table 4).

Heart and survival rates largely decreased following exposure to the Mock-GBH compared to controls (Fig 6, Table 4 and S8). However, the Roundup-WGK, Rodeo-Recommended and Rodeo-2% solutions decreased heart and survival rates to a much greater extent (Fig 1-3, Table 4, S3–5 Figs). This is despite the Mock-GBH containing comparable levels of glyphosate to Roundup-WGK and Rodeo and, in the case of Roundup-WGK, comparable levels of POEA.

These results suggest that there are other unlisted ingredients in addition to POEA in both Roundup-WGK and Rodeo that have deleterious effects on *D. magna*. The results of the Roundup-WGK and Rodeo-2% experiments suggest that the amount of unlisted ingredients in a given stock solution is proportional to the negative effects of these GBHs on *D. magna*. Although both stock solutions contain 2% glyphosate, Roundup-WGK with 98% unlisted ingredients had a much greater effect on heart and survival rates compared to Rodeo-2% with approximately 1.72% unlisted ingredients.

Our results agree with previous studies questioning the safety of the unlisted ingredients in GBHs (27,29,37,38). This is supported by our results that show greater negative impacts on heart rates and death rates of the full GBH products compared to our mock GBH solution or glyphosate alone. The 5% solution of the Roundup-WGK killed all the *magna* within 10 minutes, while the solution of 5% Mock-GBH caused a death rate of 50% within the 8-hour observation period (Table 4, S3 Fig, S8 Fig). Considering that the 5% solution of Mock-GBH contains more glyphosate by mass than the Roundup-WGK because of different salt forms of glyphosate, one would have expected that the Mock GBH would have been more harmful, based on the glyphosate alone. In comparing the other concentrations and solutions, we found similar results that indicate more harmful effects with the full GBH product compared to our solutions of the known ingredients. Solutions of glyphosate alone did not cause any death in our experiments and caused no clear effects on heart rates while higher concentrations of full GBH products Roundup-WGK and Rodeo caused increasing death and decreasing heart rates.

These results challenges the veracity of the EPA-stated safety of herbicides since they generally do not test and do not require reporting of the full ingredient makeup of commercial herbicide preparations containing glyphosate (36). This also underscores the limitations in the policy of not investigating the unlisted ingredients and total product when determining the safety of GBHs and other herbicides by regulatory and licensing agencies.

Challenges to furthering the research on the safety of GBHs as well as other herbicides/pesticides include not only the identification of the ingredients, but also their accessibility for testing. For example, in the case of glyphosate, although there are over 30 chemical vendors that sell over a dozen chemical variations, these vendors primarily sell very large quantities (1000 liter minimum order) to manufacturers of herbicides (6,10). In this investigation, we were unable to procure glyphosate formulations other than pure glyphosate. For researchers to accurately assess safety concerns, we propose three changes to the current GBH research environment:

A. Glyphosate formulations and POEA should be readily accessible in small quantities suitable for laboratory testing.
B. All unlisted ingredients in GBHs should be disclosed by herbicide manufacturers.
C. Glyphosate and POEA formulations should be disclosed by vendors to allow standardization for testing.

We hope our investigation emphasizes the need for peer-reviewed research of herbicide safety and the need to improve the transparency of product testing. This could improve the public’s confidence in government safety assessments.

Most importantly, we hope that further investigations using other organismal systems will test the listed and unlisted ingredients, as well as the full GBH product, to reveal the direct and indirect risks to human health and the environment as a result of their continually increasing use.

## Supporting information

Supplemental Data Set 1

Supplemental Data Set 2

Supplemental Data Set 3

Supplemental Data Set 4

Supplemental Data Set 5

## 5 Acknowledgements

We thank the *New Hampshire Academy of Science* for providing access to the extensive equipment of their STEM Lab to conduct our research. We are grateful for the guidance and support of Dr. Chery Whipple during the experimental phase and Elaine Faletra during the writing period of the manuscript. We especially thank Lin Chu and Dr. Zheng Duan for their technical help with data analysis and manuscript formatting.

## 7 Supporting information

**S1 Figure.**
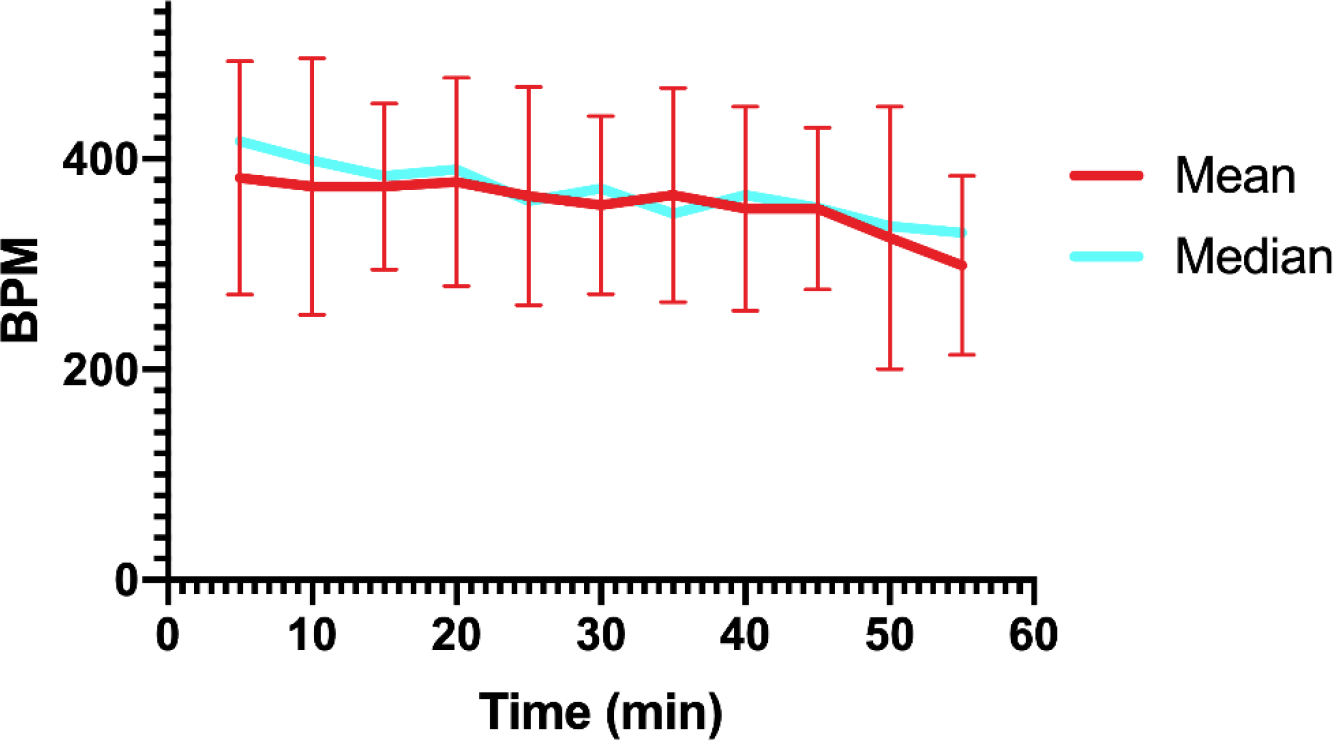
Control heart rate verification. This represents the average and median BPM of our control *D. magna* with error bars representing standard deviations. Each data point represents 25 *D. magna*. The average heart rate is shown with error bars representing standard deviation while the median heart rate is plotted without error bars. The median and mean resting heart rate in our experiments were 369 and 357 BPM respectively.

**S2 Figure.**
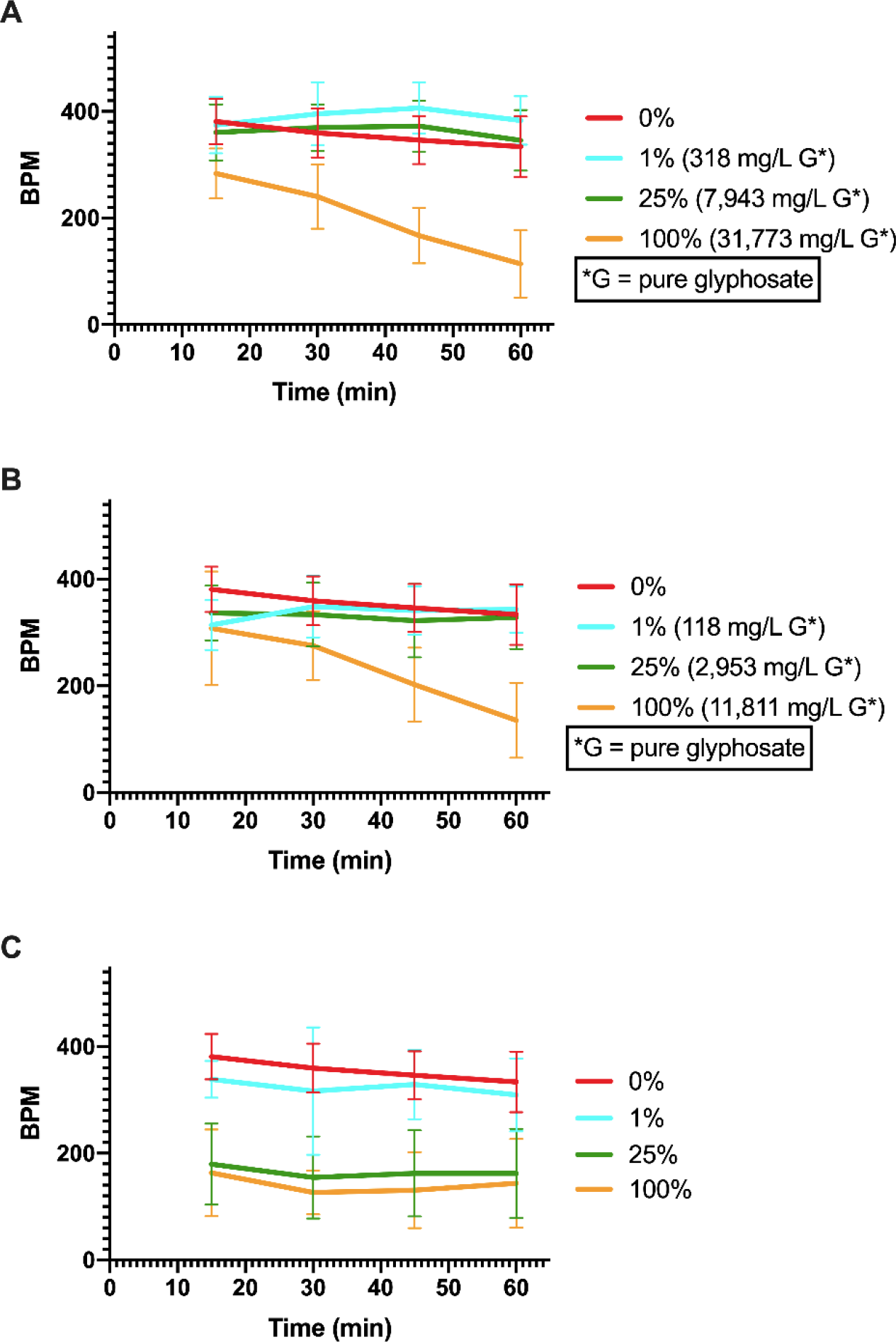
Verification of Rodeo-Recommended, Rodeo-2%, and POEA heart rate experiments. The results of the verification experiments, showing the average BPM with standard deviation of *D. magna* over exposure time in minutes. Each data point represents 10 *D. magna*. All experiments were performed at the same time and share a control though they are plotted separately for clarity. **A)** Rodeo-Recommended verification with concentrations of 1%, 25%, and 100% Rodeo-Recommended stock (100% stock contains 5.38% glyphosate). Both the 1% and 25% concentrations had little effect on the Daphnia while 100% caused heart rates to steadily drop to under 50% of the control by 60 minutes. **B)** Rodeo-2% verification with concentrations of 1%, 25%, and 100% Rodeo-2% stock (100% stock contains 2% glyphosate). Both the 1% and 25% concentrations had little effect on the daphnia while 100% caused heart rates to steadily drop to under 50% of the control by 60 minutes. This was a significant decrease compared to the control (p<0.05) **C)** POEA verification with concentrations of 1%, 25%, and 100% POEA stock (100% stock contains 1% POEA and no glyphosate). The heart rates of *D. magna* exposed to the 25% and 100% concentrations had significantly lower heart rates at less than 50% of control (p<0.05).

**S3 Figure.**
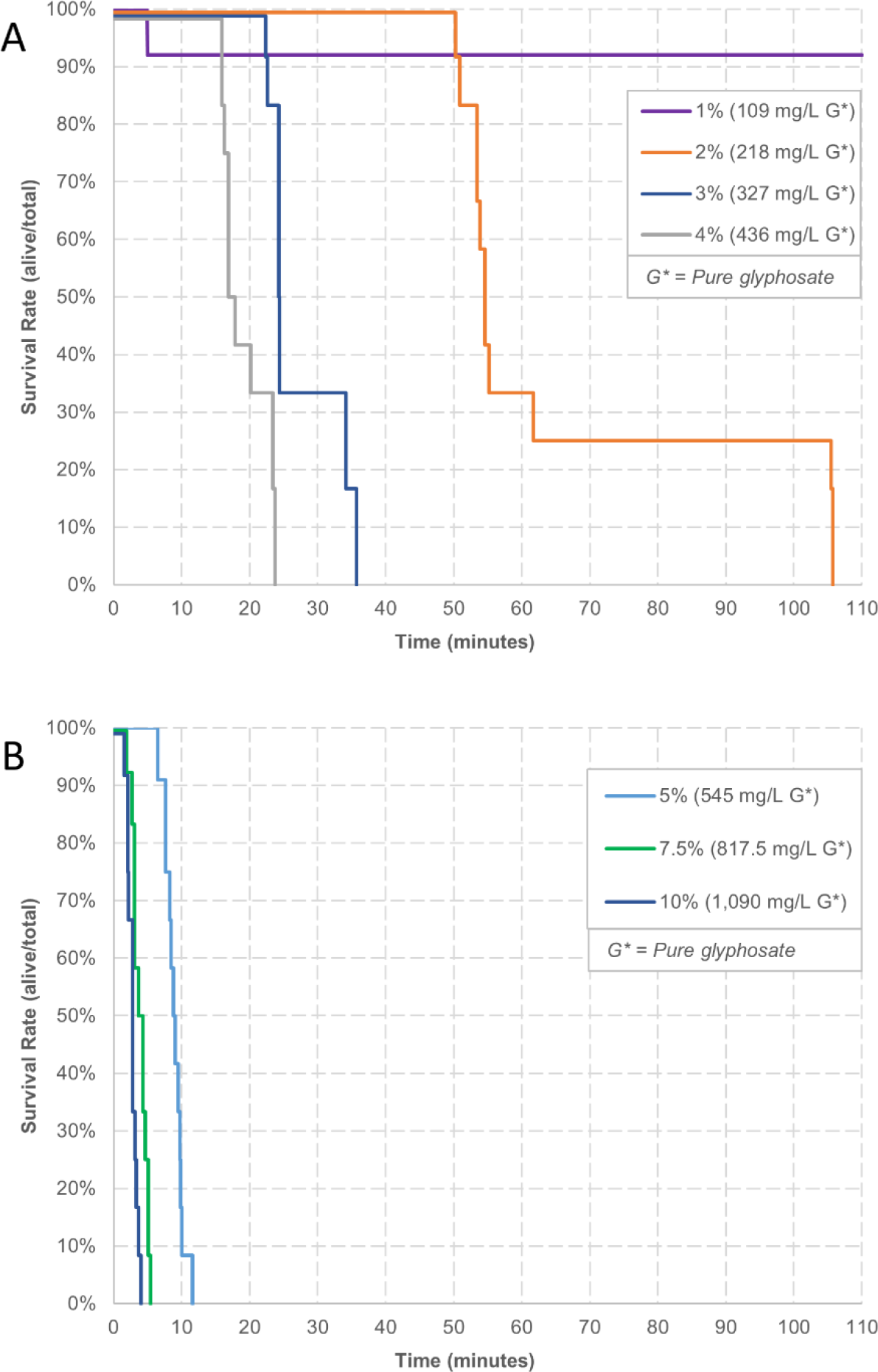
Roundup-WGK survival. This represents the survival rates of the *D. magna* exposed to Roundup-WGK dilutions over exposure time in minutes. **A)** Survival for concentrations 1%, 2%, 3%, 4% Roundup-WGK (100% stock contains 2% glyphosate). **B)** Survival in concentrations 5%, 7.5%, and 10% Roundup-WGK followed a clear dose-response pattern. Survival rates show a precipitous decline starting at 2% (A and B). Twelve *D. magna* were used for each concentration. Control group values, which had no deaths, are not shown to avoid obscuring test concentration data points. Observations were continuous over 8 hours and each death is shown by a vertical line drop at the time of death. This graph only displays the first 120 minutes of the 8-hour observational period to allow for discrimination among the groups. To avoid the colored lines representing different test groups obscuring each other, lines are slightly shifted to allow for clear visual recognition.

**S4 Figure.**
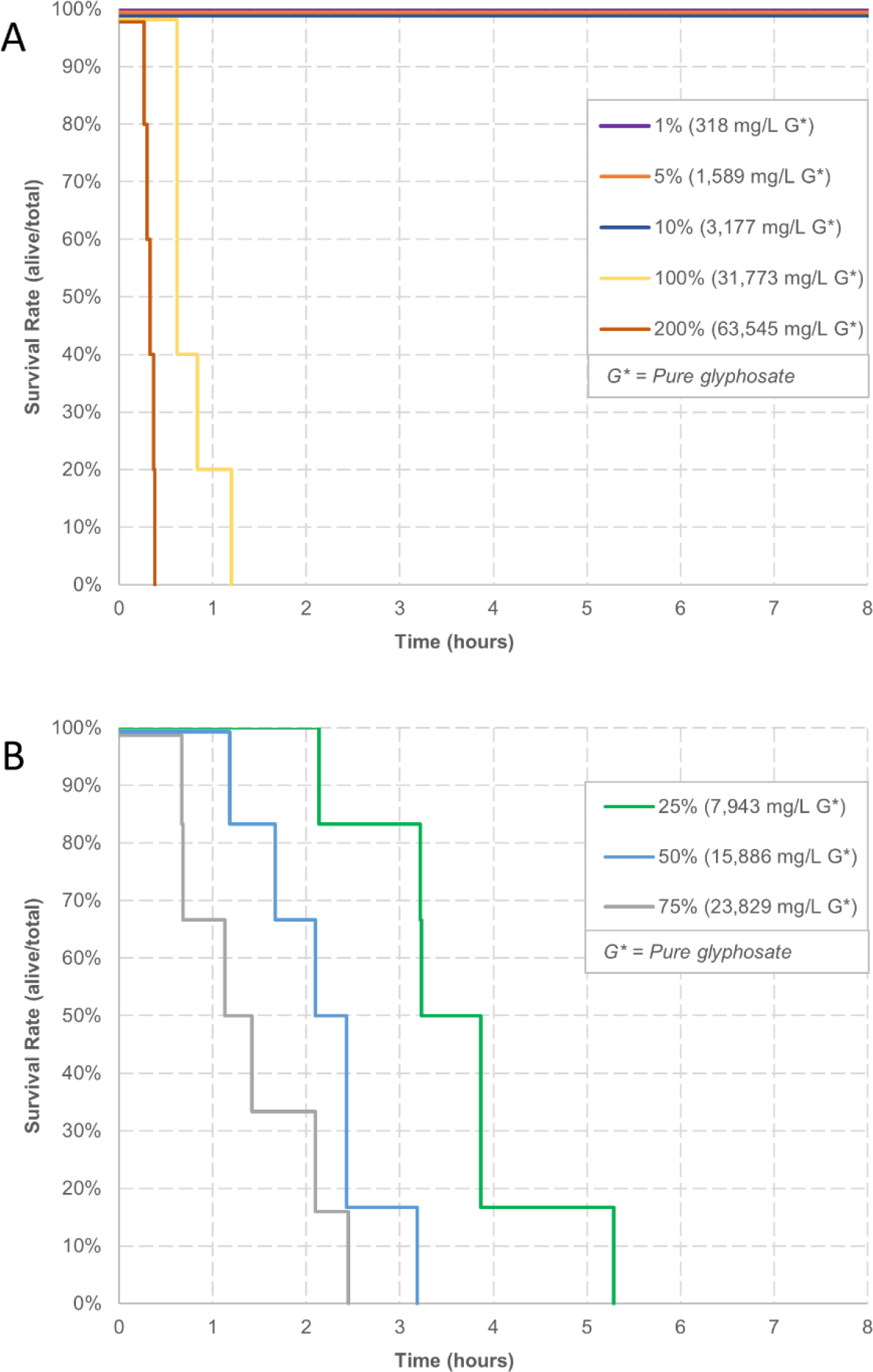
Rodeo-Recommended survival. This represents the survival rates of the *D. magna* in Rodeo-Recommended concentrations over exposure time in hours. **A)** Survival for concentrations 1%, 5%, 10%, 100%, and 200% Rodeo-Recommended stock (100% stock contains 5.38% glyphosate). **B)** Survival for concentrations 25%, 50%, and 75% Rodeo-Recommended stock, Five *D. magna* were used for each of concentrations 1%-10%, 100% and 200%; six *D. magna* were used for the remaining concentrations. Starting at 25%, death rates show a precipitous decline with a clear dose-response pattern. Control group values, which had no deaths, are not shown in graphs to avoid obscuring test concentration data points. Observations were continuous over 8 hours and each death is shown by a vertical line drop at the time of death. To avoid the colored lines representing different test groups obscuring each other, lines are slightly shifted to allow for clear visual recognition.

**S5 Figure.**
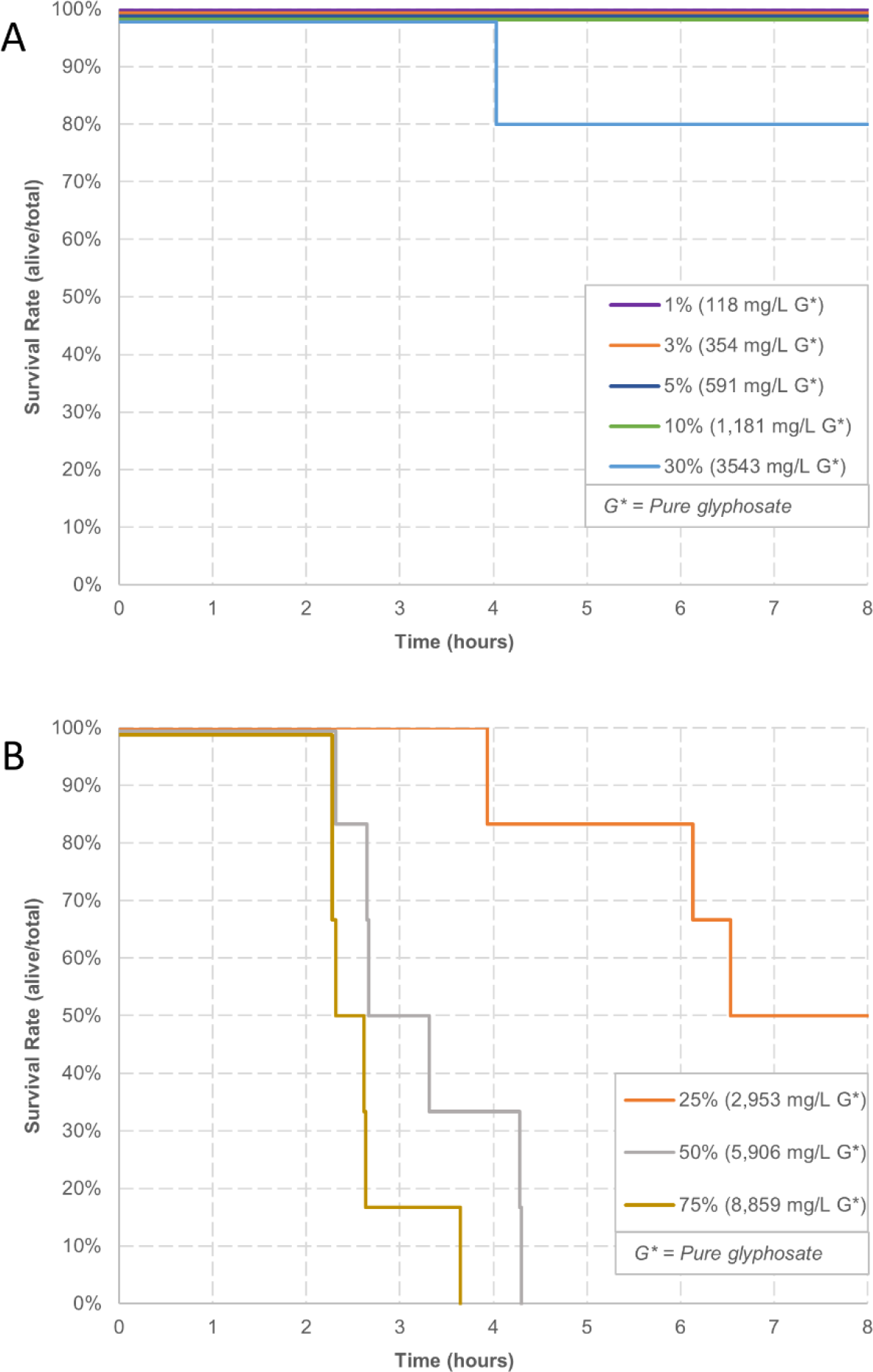
Rodeo-2% survival. This represents the survival rates of the *D. magna* over exposure time in minutes. **A)** Survival for concentrations 1%, 3%, 5%, 10%, and 30% Rodeo-2% stock (100% stock contains 2% glyphosate). B) Survival for concentrations 25%, 50%, and 75% Five *D. magna* were used for each of concentrations 1%-10% and 30%, six *D. magna* were used for the remaining concentrations. Survival rates followed a dose-response pattern starting at the 25% concentration with the two highest concentrations showing a precipitous decline within 5 hours. Control group values, which had no deaths, are not shown in graphs in order to avoid obscuring test concentration data points. Observations were continuous over 8 hours and each death is shown by a vertical line drop at the time of death. To avoid the colored lines representing different test groups obscuring each other, lines are slightly shifted to allow for clear visual recognition.

**S6 Figure.**
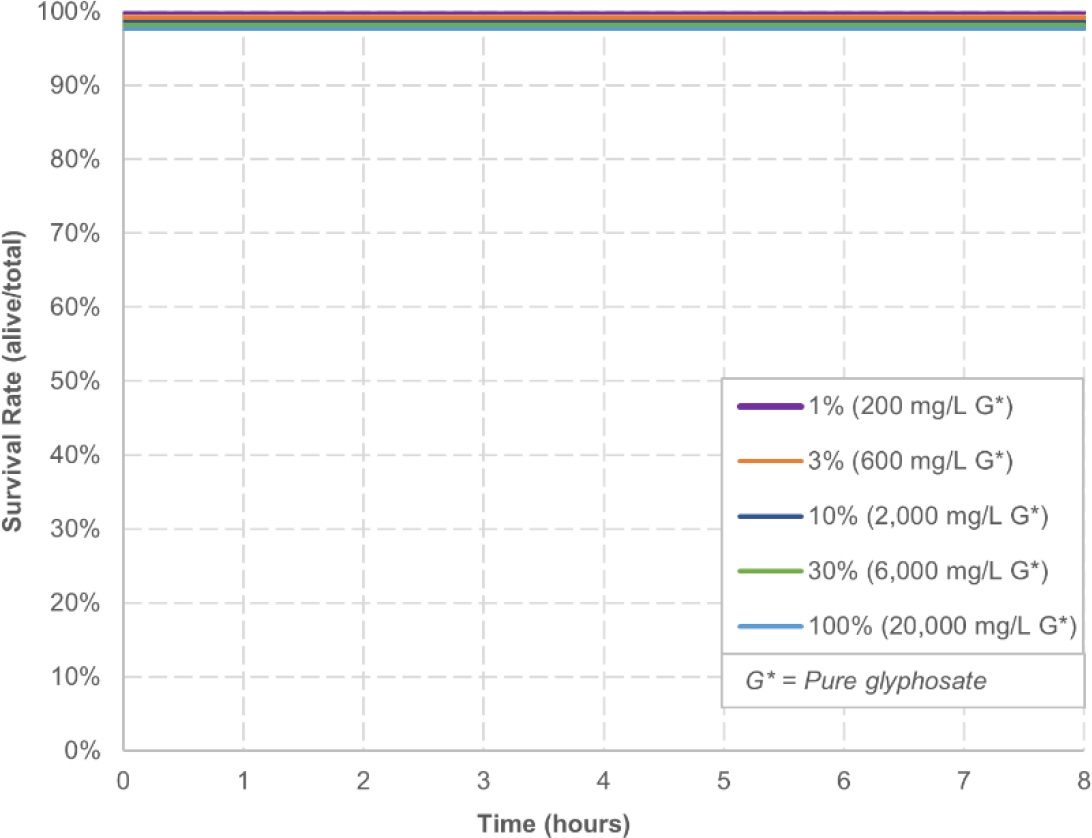
Glyphosate survival. This graph shows the survival rates of *D. magna* over an 8-hour exposure time for 1%, 3%, 10%, 30%, and 100% concentrations of the glyphosate stock solution. Six *D. magna* were used for each concentration. No *Daphnia* died during the observation times for any concentrations. Control group values, which had no deaths, are not shown in graphs to avoid obscuring test concentration data points. Observations were continuous over 8 hours. To avoid the colored lines representing different test groups obscuring each other, lines are slightly shifted to allow for clear visual recognition.

**S7 Figure.**
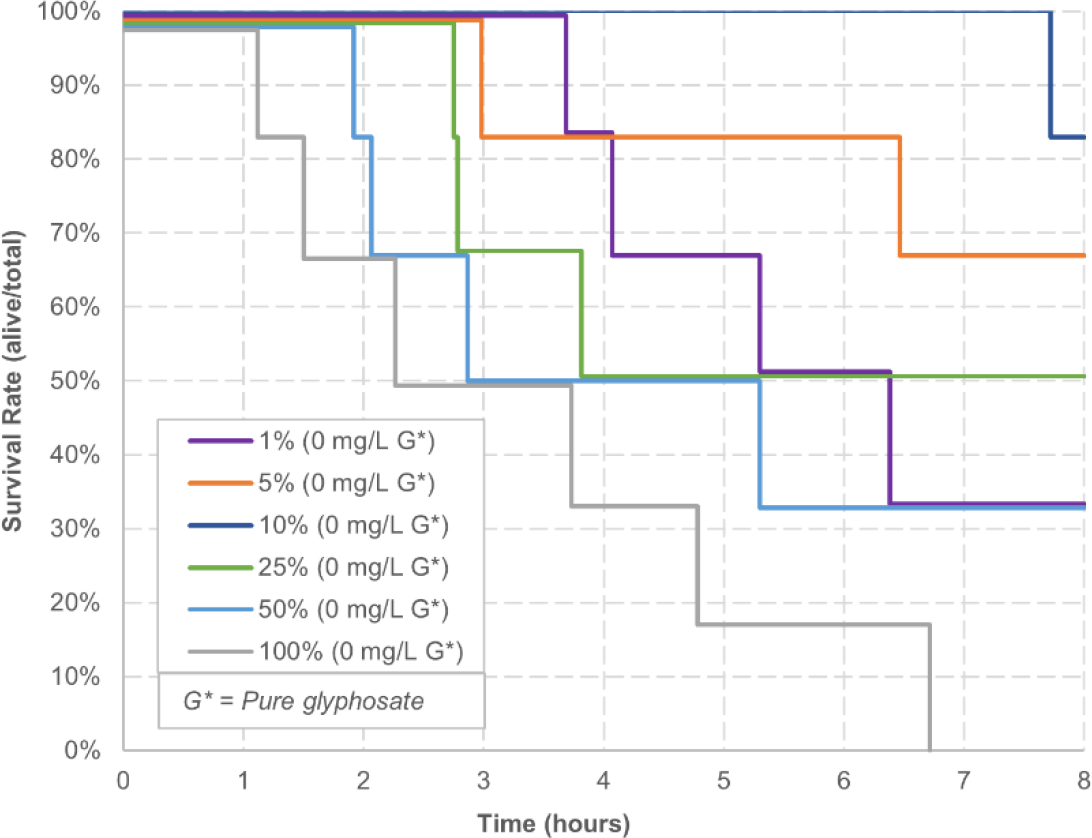
POEA survival. This represents the survival rates of the *D. magna* over exposure time in hours for concentrations 1%, 5%, 10%, 25%, 50%, and 100% POEA (100% stock contains 1% POEA and no glyphosate). Six *D. magna* were used for each concentration. Survival rates follow a dose-response pattern for the three highest concentrations dropping to 50% survival within the first 4 hours. The 1%, 5%, and 10% concentrations did not follow a dose-response pattern. Control group values, which had no deaths, are not shown in graphs to avoid obscuring test concentration data points. Observations were continuous over 8 hours and each death is shown by a vertical line drop at the time of death. To avoid the colored lines representing different test groups obscuring each other, lines are slightly shifted to allow for clear visual recognition.

**S8 Figure.**
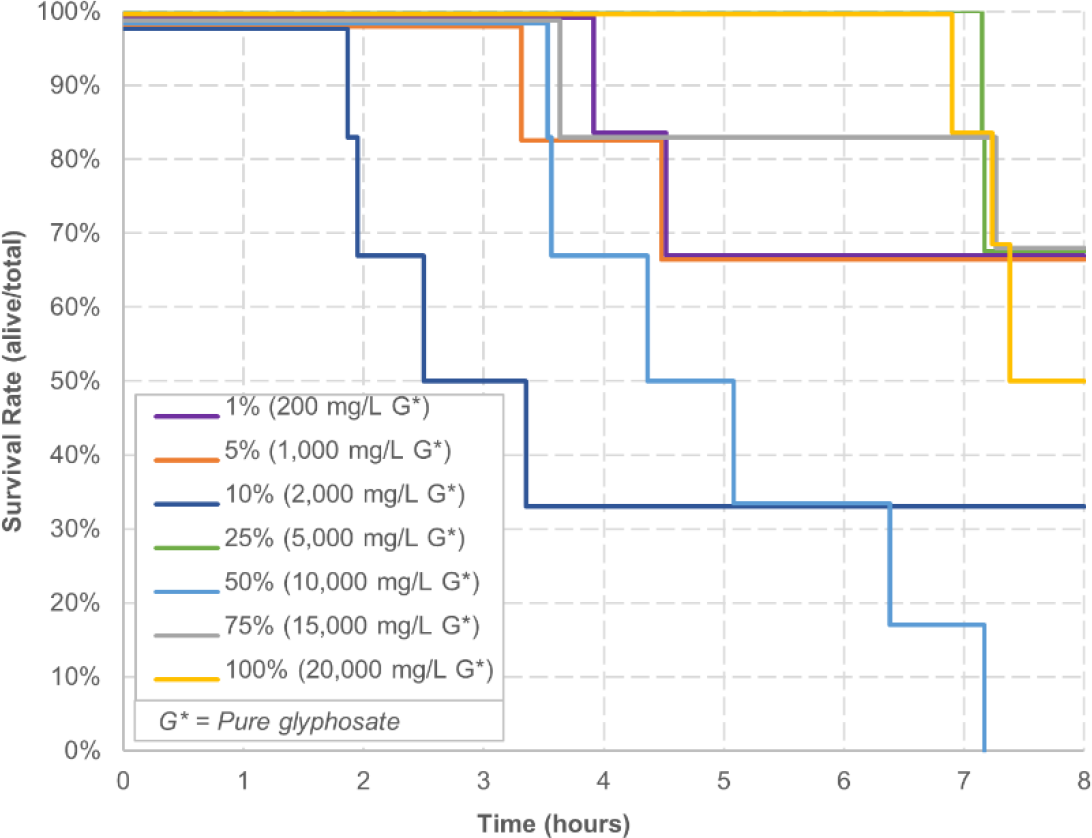
Mock-GBH survival. This represents the survival rates of the *D. magna* over exposure time in hours for concentrations 1%, 5%, 10%, 25%, 50%, 75%, and 100% Mock-GBH (100% stock contains 1% POEA and 2% glyphosate). Five to six *D. magna* were used for each concentration. Survival rates do not follow a dose-response pattern. Control group values, which had no deaths, are not shown in graphs to avoid obscuring test concentration data points. Observations were continuous over 8 hours and each death is shown by a vertical line drop at the time of death. To avoid the colored lines representing different test groups obscuring each other, lines are slightly shifted to allow for clear visual recognition.

**S1 Dataset. Statistical results for heart rate experiments**. Statistical analysis of the results was performed by applying a Kruskal-Wallis ANOVA test with Dunn’s Multiple Comparison post-test using GraphPad Prism.

**S2 Dataset. Raw data for heart rate experiments.**

**S3 Dataset. Raw data for heart rate verification experiments of Rodeo-Recommended, Rodeo-2% and POEA**.

**S4 Dataset. Raw data for survival experiments.**

**S5 Dataset. Consolidated control heart rate data.**

